# Inference and prediction for stochastic models of biological populations undergoing migration and proliferation

**DOI:** 10.1101/2025.05.25.656057

**Authors:** Matthew J. Simpson, Michael J. Plank

## Abstract

Parameter inference is a critical step in the process of interpreting biological data using mathematical models. Inference provides a means of deriving quantitative, mechanistic insights from sparse, noisy data. While methods for parameter inference, parameter identifiability, and model prediction are well-developed for deterministic continuum models, working with biological applications often requires stochastic modelling approaches to capture inherent variability and randomness that can be prominent in biological measurements and data. Random walk models are especially useful for capturing spatiotemporal processes, such as ecological population dynamics, molecular transport phenomena, and collective behaviour associated with multicellular phenomena. This review focuses on parameter inference, identifiability analysis, and model prediction for a suite of biologically-inspired, stochastic agent-based models relevant to animal dispersal and populations of biological cells. With a particular emphasis on model prediction, we highlight roles for numerical optimisation and automatic differentiation. Open-source Julia code is provided to support scientific reproducibility. We encourage readers to use this code directly or adapt it to suit their interests and applications.

## 1 Introduction

Mathematical models are an increasingly ubiquitous tool in the life sciences. Frequently, models are built from domain-specific knowledge about the system of interest, and typically contain parameters whose values are *a priori* unknown. Parameter inference refers to statistical methods by which these parameters may be inferred from available data, which are often noisy and incomplete, enabling us to draw conclusions about biological mechanisms or to make predictions about future observations. Whether we are interested in studying disease progression [1], ecological population dynamics [2], cell fate decisions [3], or gene regulation [4], mathematical and computational methods are routinely used to estimate parameters, test hypotheses, and quantify uncertainty [5]. Inference techniques include classical statistical approaches, Bayesian methods, and machine learning, enabling us to integrate diverse data sources, explore hidden patterns, to test and validate mechanistic models.

Continuum models, which describe systems in terms of continuous variables (e.g. density, temperature, displacement) are widely used in classical mathematical modelling applications such as fluid dynamics [6] and material science [7]. Inference for continuum models is well-established across many fields of science and engineering, supported by established foundations in both Bayesian and frequentist methodologies [8–11]. Frequentist approaches, including maximum likelihood estimation and hypothesis testing, have long provided tools for parameter estimation, model validation and model selection [8, 11]. Meanwhile, Bayesian inference incorporates prior knowledge, uncertainty quantification, and performing model selection [9, 10, 12]. The maturity of inference techniques for continuum models has enabled reliable calibration of mathematical models based on ordinary differential equations (ODEs) and partial differential equations (PDEs) across a broad range of applications.

Robust inference in life sciences applications is particularly critical due to the inherent challenges posed by non-standard data collection protocols and limited knowledge of the under lying mechanistic principles, especially when compared to fields like physics and engineering. Biological data are often sparse, noisy, and collected under varying experimental and field conditions. These features can complicate the tasks of parameter estimation, parameter identifiability analysis, and model prediction. Unlike in physics and engineering, where governing equations are typically well-known and able to be derived from first principles, research in life sciences frequently involves phenomenological models that act as coarse approximations of the true biological processes. As a result, careful inference is essential not only for estimating parameters but also for assessing model identifiability, validating assumptions, appropriately capturing uncertainty, and guiding model refinement.

As data collection protocols and imaging technologies continue to advance, the availability of high-resolution spatiotemporal data is transforming the way biological systems are modelled and analysed [13–19]. Detailed biological datasets capture variability and noise that are often masked in coarse-grained continuum modelling frameworks. This motivates the development and application of stochastic models that more accurately reflect the inherent randomness of biological measurements. In particular, random walk models and related stochastic frameworks have gained prominence for describing a broad range of phenomena including animal movement ecology, multicellular phenomena, molecular diffusion, and intracellular transport [20, 21]. Within the theoretical biology literature, the task of drawing inferences from such models is predominantly approached through Bayesian methods, which are well-suited for handling uncertainty, integrating prior knowledge, and dealing with complex model structures. However, this Bayesian dominance often leads to the undervaluation and under-utilisation of frequentist approaches, which can offer complementary strengths, such as computational efficiency, established theoretical guarantees, and interpretability [22–26].

This review examines the interrelated topics of inference, identifiability, and prediction for stochastic random walk models, and their generalisations. We are particularly interested in applications in population biology, where populations are composed of individuals that undergo some form of random movement and reproduction. Therefore, these models and techniques are well-suited to understanding problems in animal dispersal and collective motion of populations of biological cells. Multi-agent, simulation-based random walk models provide a natural and flexible framework for capturing movement, reproduction and spatial dynamics in these systems, especially where stochasticity plays a central role. We will consider both Bayesian and frequentist approaches to inference, highlighting their respective strengths and limitations in the context of biological data. Special attention will be given to using surrogate models to approximate complex likelihoods, as well as using automatic differentiation to streamline sampling for model prediction. All code required to replicate our results are available as open source Julia code on GitHub, and we encourage readers to use this code directly or adapt it to suit their interests and applications.

## 2 Motivation

In this Section, we discuss a simple experimental framework that will motivate our-high fidelity, computationally expensive discrete random walk model that aims to describe key features of the experimental mechanisms and observations.

### 2.1 Experimental observations

The initial experimental motivation for our stochastic simulations is a barrier assay experiment describing the spatial expansion of a circular monolayer of cells, such as the outward circular expansion of a population of fibroblast cells shown in Figure 1(a)–(b) [27]. In this experiment [28, 29], cells are placed into a circular barrier and some time is allowed for individual cells to attach to the cell culture plate. Non-viable cells, and those cells that remain unattached to the plate are removed [27] and the barrier is lifted to initiate the experiment at *t* = 0. The subsequent outward radial expansion of the remaining attached population is imaged, such as the image presented at *t* = 72 hours in Figure 1(b). In this particular experiment, all cells were treated with a chemical inhibitor to prevent proliferation and there was no observed cell death [27]. Images captured during these experiments clearly show the role of stochasticity since random locations of individual cells within the expanding population are visible and can be measured. This kind of experiment motivates us to understand how we can infer macroscopic transport coefficients (e.g., cell diffusivity) and quantify the density of viable cells that have attached to the surface of the cell culture plate at *t* = 0 as the barrier is lifted.

**Figure 1:**
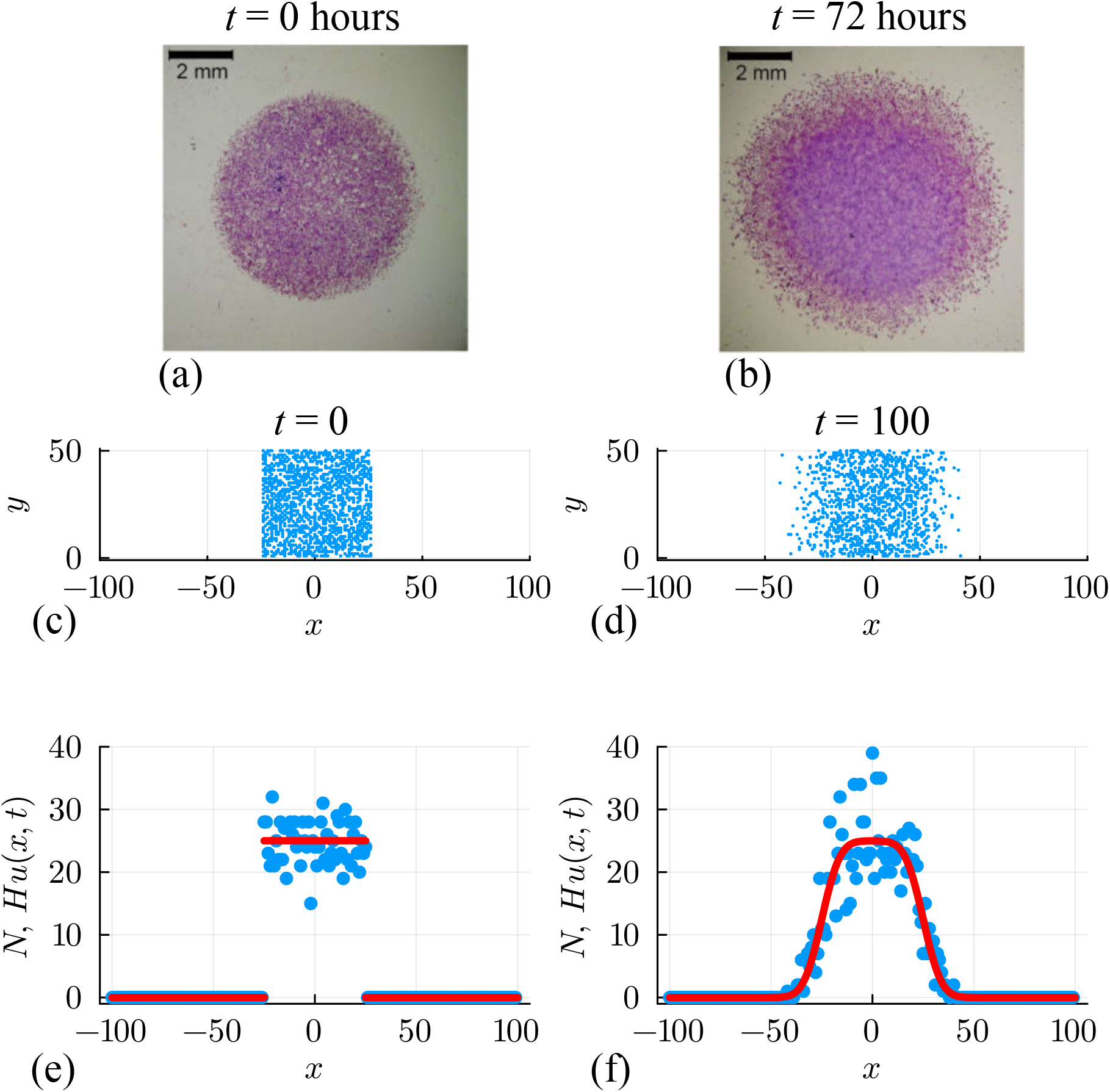
(a)–(b) Barrier assay experiment showing: (a) the initial placement of a population of fibroblast cells within a confined circular region of diameter 6 mm; and, (b) the outward spreading of individual cells after 72 hours [27]. (c)–(d) Unbiased random walk simulation on a square lattice with Δ = *τ* = 1, *H* = 50 and *L* = 100. The initial placement of agents in a confined region is shown in (c), and the subsequent outward spreading of agents after 100 time steps is shown in (d). Initially, sites in the region −25 ≤ *x* ≤ 25 are occupied by, at most, a single agent with probability *U* = 0.5, and then random agent migration takes place with *P* = 1. (e)–(f) Numbers of agents per column at *t* = 0 and *t* = 100, respectively. Noisy count data (blue dots) are superimposed with *Hu*(*x, t*) (solid red curve), where *u*(*x, t*) is the solution of Equation (2) with *D* = *P/*4.

### 2.2 Discrete model

We first implement a classical non-interacting unbiased random walk model to illustrate some fundamental concepts [20, 30–32]. Later we will extend this fundamental modelling framework to include directional bias, proliferation of agents, and crowding effects. To commence, we consider a lattice-based random walk model on a two-dimensional square lattice with lattice spacing Δ. Each lattice site is indexed in the usual way (*i, j*), so that each site is associated with location (*x*_*i*_, *y*_*j*_), for *i* = 1, 2, 3, …, *I* and *j* = 1, 2, 3, …, *J*. Simulations are performed on a lattice of height *H* and width *W*, so that 0 ≤ *y* ≤ *H* and −*L* ≤ *x* ≤ *L*, where *W* = 2*L*. In this case we have *x*_*i*_ = −*L* + (*i* − 1)Δ for *i* = 1, 2, 3, …, *I* and *y*_*j*_ = (*j* − 1)Δ for *j* = 1, 2, 3, …, *J*. In this standard model, each lattice site can be occupied by multiple agents, and agents undergo an unbiased random walk within a discrete time stepping framework where each time step has duration *τ* [20, 30–32]. To evolve the system from time *t* to time *t*+*τ*, we consider a random sequential update method [33] which, for a population of *Q* agents, involves selecting *Q* agents independently at random, one at a time, with replacement. Each selected agent moves with probability *P* ∈ [0, 1]. If an agent attempts to move, the target site is chosen randomly from the four nearest neighbour sites. To maintain generality, we always work with dimensionless simulations by setting Δ = *τ* = 1, noting that these simulations can be re-dimensioned to match any dimensional lattice spacing and time step [34]. We will demonstrate the relationship between dimensional experimental data and dimensionless simulations through a Case Study in Section 5.

In this preliminary simulation, we attempt to capture key features of the cell biology experiments in Figure 1, namely we consider a population of cells are initially placed into a confined region, and then we use the discrete model to capture snapshots and data from the spreading population for *t >* 0 [27]. At the beginning of the experiment, *t* = 0, the confined population is shown in Figure 1(a). As the experiment proceeds the population of cells spreads symmetrically outward over time, as illustrated in Figure 1(b). To lay out our analysis as straightforward as possible, we consider simulations performed on a relatively long and narrow lattice with *H* = 50 and *W* = 200 so that −100 ≤ *x* ≤ 100 and 0 ≤ *y* ≤ 50. Zero flux boundary conditions are applied along all boundaries (i.e., if an agent attempts to move in a direction that would take it off the lattice, that potential move is aborted). Simulations are initialised by randomly occupying each site within the region where |*x*| ≤ 25 by a single agent with probability *U* = 0.5, leading to the distribution of agents in Figure 1(c). A single simulation is performed by allowing the system to evolve over 100 time steps with *P* = 1, with the output of the simulation leading to a population that spreads symmetrically in the positive and negative *x*-directions, as in Figure 1(d). Since there is no variation in the initial macroscopic density in the *y*-direction, the resulting distribution of agents remains, on average, independent of vertical location for this choice of initial conditions and boundary conditions [34]. This kind of experimental design is both mathematically convenient, and also used in practical cell biology experiments since it is simpler to characterise the spatiotemporal spreading of the population in terms of just one spatial coordinate [35–39]. While it is not immediately obvious from the visualisation in Figure 1(c)-(d), the initialisation of the simulation means that all lattice sites at *t* = 0 either contain zero or, at most, a single agent. In contrast, at subsequent times (e.g. as in Figure 1(d)), some lattice sites can be occupied by more than one agent since this is a non-interacting random walk model where there is no limit on the number of agents per site and we are not modelling any kind of crowding effect [21].

To quantify the simulations we let *n*_*i,j*_ (*t*) ∈ [0, ∞) denote the integer number of agents at site (*i, j*) at time *t*. Since the average occupancy of lattice sites is independent of vertical location, we quantify the distribution by counting the number of agents per column,

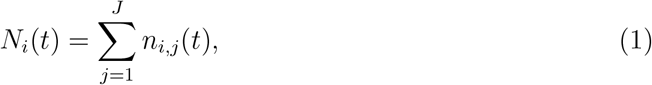

so that *N*_*i*_(*t*) ∈ [0, ∞) denotes the integer number of agents in the *i*th column at time *t*. For notational simplicity we will sometimes drop the explicit reference to the independent variables by referring to *N*_*i*_(*t*) as either *N*_*i*_ or *N*, as appropriate. This number *N* is plotted in Figure 1(e)–(f) for the simulation results at *t* = 0 and *t* = 100, respectively. In Figure 1(e) we see that we have *N*_*i*_ = 0 for sites where |*x*| *>* 25, as expected. For sites with |*x*| ≤ 25 we have noisy count data where our plot of *N*_*i*_involves large fluctuations in this region. Since these lattice sites are occupied with probability *U* = 0.5, the expected number of agents is *H × U* = 25. However, in our example with a lattice of finite height *H* = 50, some columns at *t* = 0 contain as many as *N*_*i*_ = 36 agents, whereas others contain as few as *N*_*i*_ = 17. This noise is an important feature of real experimental data, which is a key motivation for the use of a stochastic model. To illustrate the importance of these fluctuations we superimpose _the expected number of agents per column, 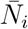_ for *i* = 1, 2, 3, …, *I*, in Figure 1(e). At *t* = 0 _we have 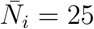 for |*x*| ≤ 25 and 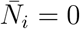_. Superimposing plots of *N*_*i* and 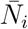_ in this way makes it clear that the data from the stochastic random walk model leads to large fluctuations, and in this review we address the question of how to meaningfully incorporate these large fluctuations using a combination of surrogate models and noise models, as we will now begin to explore.

### 2.3 Surrogate model

The mean behaviour of the discrete model is related to a PDE that describes the average density of agents as a function of continuous position and time [20, 21, 31, 32],

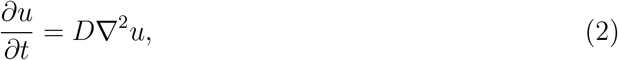

Where 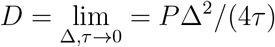, such that *u*(*x, y, t*) ≥ 0 denotes the average density of agents at location (*x, y*) at time *t*. This continuum limit is valid in the constrained limit as Δ → 0 and *τ* → 0 with the ratio Δ^2^*/τ* held constant [31]. Since our simulations do not involve macroscopic density gradients in the vertical direction, *∂u/∂y* vanishes and this PDE simplifies to

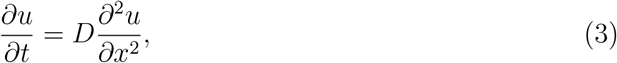

for the density *u*(*x, t*) ≥ 0, with initial condition

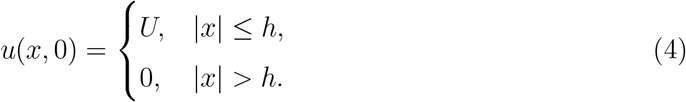

Since *u*(*x, t*) ≥ 0 represents the mean agent density, for our simulations the product *H × u*(*x, t*) is the continuous analogue of *N*_*i*_ obtained from the discrete simulations. With our choice of initial conditions, and simulations of an intermediate time scale where agents do not reach the vertical boundaries at *x* = *± L*, we can approximate the solution of Equation (3) with the appropriate infinite domain solution [40]

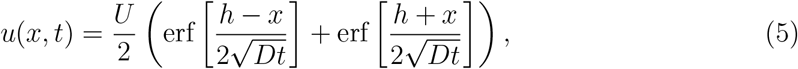

where 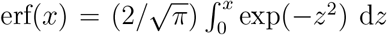 is the error function [41]. Evaluating this solution with *h* = 25 and *U* = 0.5 to match the initial condition, and *D* = 0.25 to match the jump probability, we plot the quantity *Hu*(*x, t*) at *t* = 100 in Figure 1(f) to illustrate that the solution of the PDE model provides a good prediction of the evolution of the average number of agents per column in the discrete simulation. Clearly the PDE solution does not capture the fluctuations present in the discrete simulation data, and much of this review will be devoted to exploring different ways of quantifying and capturing these fluctuations using an appropriate choice of noise model.

## 3 Inference

We will now work through a number of computational exercises aimed at using discrete count data in Figure 1(f) to infer parameters relevant to the solution of the mathematical model. As we work through various problems, we will introduce different details into the discrete model and explore the consequences of increasing the complexity of the inference problem. Open access software that can be used to replicate each case we consider is available on GitHub.

### 3.1 Approximate Bayesian Computation: Estimation and prediction

We now consider the problem of using the discrete data in Figure 1(f) to estimate values of the parameters *θ* = (*U, D*)^⊤^. In other words, we aim to take the noisy output of the discrete simulation, and estimate values of *U* and *D* that best explain this noisy data under the random walk model. Throughout this document we will refer to the data using the vector *y*^obs^, which in this case is a vector of length 201 so that the *i*th element is given by *N*_*i*_ as plotted in Figure 1(f). In the first instance we consider a Bayesian setting in which the parameters *θ* = (*U, D*)^⊤^ are treated as random variables, and the uncertainty in *θ* is updated using simulation data [42–45]. Simulation data are generated by randomly sampling values of *θ* = (*U, D*)^⊤^ from a uniform prior distribution within the region 0.05 ≤ *D* ≤ 0.40 and 0.30 ≤ *U* ≤ 0.70 since we know in advance that this interval contains the true parameter values used to generate the synthetic data. For the *j*th random sample *θ*_*j*_, we initialise the discrete model and perform a discrete simulation over 100 time steps to record the resulting count data *N*^(*j*)^, and we compute a discrepancy measure for the *j*th parameter sample

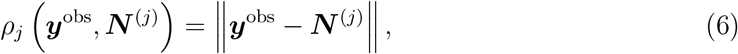

which we record as, *ε*_*j*_ = *ρ*_*j*_. To implement the algorithm we repeat this process until we obtain 10^5^samples of 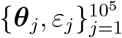. Samples are sorted in order of increasing *ε*, and we retain a small proportion of samples so that we focus on parameter combinations that provide a good match to the data. In this case we retain the top 1% of samples as posterior samples, and we checked (results not shown) that the key properties of the posterior distribution are relatively insensitive to this choice of tolerance [46]. We also checked that our ABC rejection results are insensitive to widening the prior distribution of *D* and *U*, noting that widening the prior does not have a noticeable impact on the results other than significantly slowing down the computational procedure. We encourage readers to download the Jupyter notebooks from GitHub and explore the consequences of working with different prior distributions and varying the tolerance used to construct the posterior samples.

Results of this elementary algorithm are presented in Figure 2(a), where an unsmoothed histogram of posterior samples is presented. Taking the mean of the posterior samples gives *θ* = (0.489, 0.254)^⊤^, which is relatively close to the expected result, *θ* = (0.500, 0.250)^⊤^. The shape of the histogram indicates that the posterior samples for *U* are relatively narrowly distributed about the mean, whereas the posterior samples of *D* are not. The marginal univariate distributions for *D* and *U*, also presented in Figure 2(a), show this more clearly with the posterior distribution of *U* relatively narrow about the mean, whereas the posterior distribution of *D* is relatively flat and the credible interval is not very different from the width of the prior distribution. While there are many possibilities for improving the performance of this simple approach (e.g. by exploring different summary statistics or different sampling methods) we will now explore the implications of repeating the process using a surrogate model.

**Figure 2:**
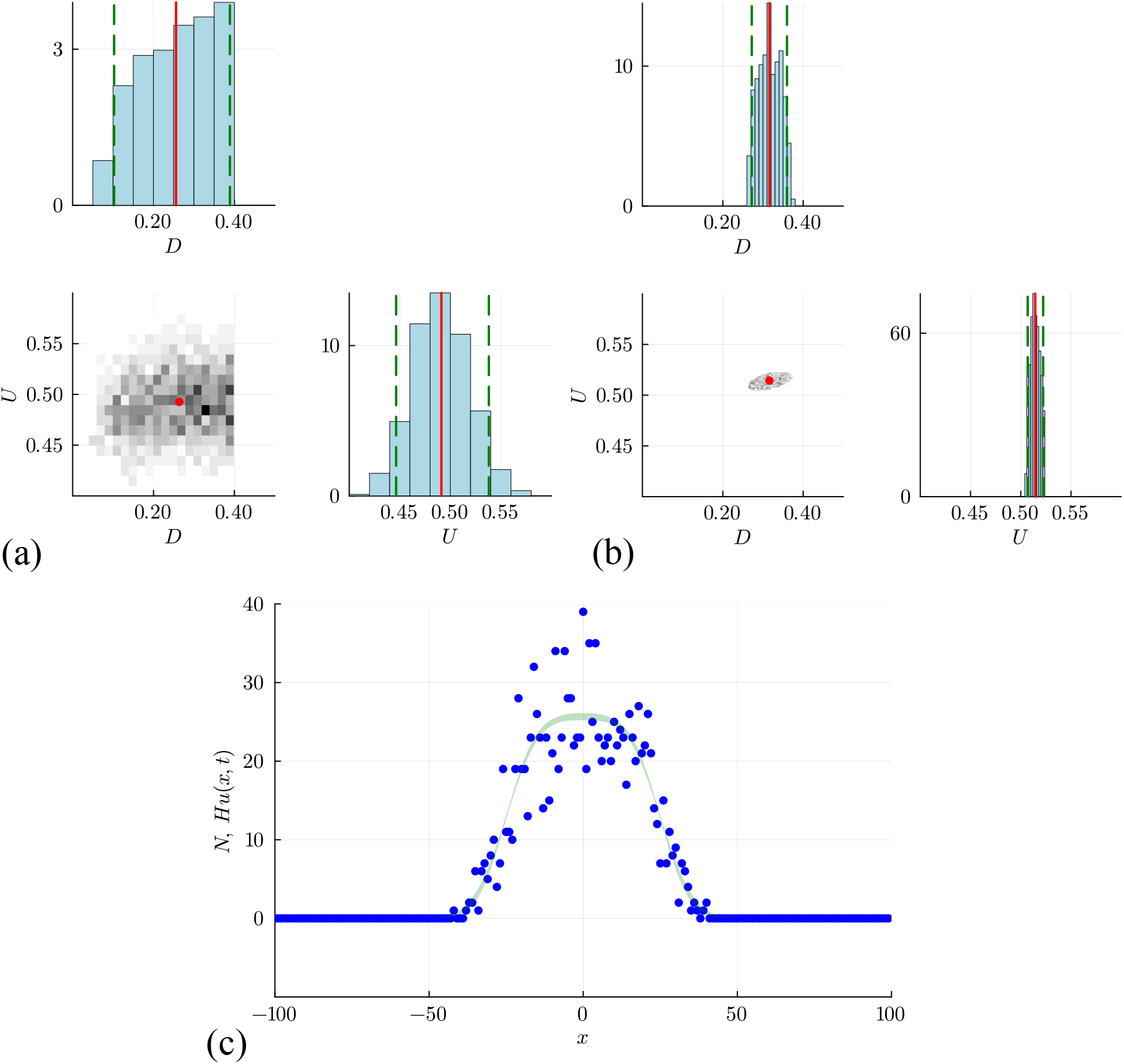
ABC rejection results based on: (a) the discrete simulation model, leading to point estimates of *D* = 0.254 [0.091, 0.394] and *U* = 0.489 [0.440, 0.548]; and, (b) the surrogate PDE model, leading to estimates of *D* = 0.317 [0.269, 0.367] and *U* = 0.514 [0.506, 0.523]. Point estimates correspond to the mean of the posterior samples, and the credible interval is obtained by computing the 0.025 and 0.975 quantiles of the posterior samples. In (c) we superimpose the data, *N*_*i*_, and the credible interval for *N* obtained by taking solving the PDE model for each of the *M* = 1000 parameter samples and plotting the point-wise union of these *M* solutions, *Hu*_*m*_(*x, t*) for *m* = 1, 2, 3, …, *M*, where each solution is evaluated on a uniform discretisation of the domain −100 ≤ *x* ≤ 100, discretised using 201 mesh points.

Results in Figure 2(b) summarise the outcome of the same ABC rejection algorithm, using the same data as before, except now we work with the solution of the continuum limit model, Equation (5). As discussed before, the solution of this continuum limit model gives *u*(*x, t*), and taking the product *Hu*(*x, t*) gives us the expected number of agents in the column at location *x* under the solution parameterised with *θ* = (*U, D*)^⊤^. The outcome of the *j*th model simulation is a vector 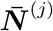, whose *i*th element is given by *Hu*(*x*_*i*_, *t*), where *x*_*i*_is the horizontal location of the *i*th column in the lattice. For each 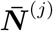 we compute *ε*_*j*_ = *ρ*_*j*_ and retain 1% of 10^5^random samples with the smallest discrepancy and then generate the histogram of posterior samples as shown in Figure 2(b). The posterior mean is *θ* = (0.514, 0.317)^⊤^. Compared to the distributions in Figure 2(a), the posterior samples in Figure 2(b) obtained using the surrogate PDE are much more narrowly distributed about the mean. In this case, however, the credible interval for *D* does not contain the expected true value, *D* = 0.25. This could be due to some bias in the estimator, or it could simply be the result of the extremely variable data, as previously discussed with reference to the data in Figure 1(f). Since we have obtained reasonably precise (but potentially inaccurate) estimates of *U* and *D*, we can solve the process model, Equation (5), for each of the *M* = 1000 retained parameter samples, giving a series of *M* solutions, *u*_*m*_(*x, t*) for *m* = 1, 2, 3, …, *M*. With these *M* solutions, we construct a credible interval for *N* by uniformly discretising the domain, −100 ≤ *x* ≤ 100 using 201 equally-spaced mesh points. At the *i*th mesh point we form a credible interval for *N* by finding the maximum and minimum value of *Hu*_*m*_(*x*_*i*_, *t*), for *m* = 1, 2, 3, …, *M* (see Figure 2(c)). While this relatively narrow credible interval for *N* captures the mean trends in the data reasonably well, it does not capture the fluctuations in the data.

Before moving on to a different approach for estimation, we note there are several options available for reporting point estimates from posterior samples, such as those in Figure 2(a)–(b). For simplicity, in this work we report the posterior mean. For example, for the posterior samples in Figure 2(b) the posterior mean for the initial occupancy gives a point estimate of *U* = 0.514306, given here with six decimal places. Alternatively, it could be useful to report the maximum a posteriori (MAP) estimate, giving *U* = 0.514445 for this data. Here, we see that these point estimates are not very different which is consistent with our observation that the posterior distribution is unimodal and approximately symmetric. Therefore, for simplicity, we report the posterior mean, noting that the posterior mean is mathematically related to working with an additive Gaussian noise model [47, 48] that we will now introduce.

### 3.2 Likelihood-based sampling: Estimation and prediction

We will now turn to a likelihood-based sampling approach for the same inference question of estimating the parameters *θ* = (*U, D*)^⊤^ that best explain the data in Figure 1(f) using the solution of the relevant surrogate model, Equation (5). Unlike the likelihood-free ABC rejection approaches outlined in Section 3.1, here we introduce an observation model, also known as a noise model or measurement error model [49], that relates the solution of the mathematical model at the *i*th lattice site, *Hu*(*x*_*i*_, *t*), to the observed count data, *N*_*i*_, by assuming that the data are normally distributed about the solution of the mathematical model with constant variance *σ*^2^. This assumption can be expressed mathematically as

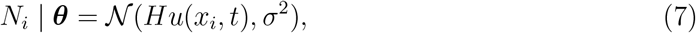

at each column of the lattice, *i* = 1, 2, 3, …, *I*. At this point we have provided no justification to support the standard assumption that the data are normally distributed about the solution of the model [49–53]. For the moment we will proceed under this assumption, and comment on the suitability of this standard approach later. Before proceeding it is worth noting that this assumption introduces another unknown parameter *σ*^2^, and we will estimate this additional parameter from the data. This means that we will work with the expanded vector of parameters *θ* = (*U, D, σ*)^⊤^. One advantage of introducing the measurement error model is that we now have an objective way of quantifying how close the solution of the mathematical model is to the data, without the need to specify a distance function like we did in Section 3.1. Here we measure the closeness of the data and the model solution via the probability density associated with the normal distribution. For a series of measurements at position *x*_*i*_for *i* = 1, 2, 3, …, *I*, invoking a standard independence assumption motivates us to take the product of *I* densities to give the likelihood of observations given the solution of the mathematical model parameterised by *θ*,

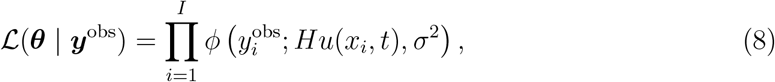

where *ϕ*(*x*; *µ, σ*^2^) denotes the probability density function of the normal distribution with mean *µ* and constant variance *σ*^2^[50, 51]. Under this approach, parameters of the mathematical model that lead to an improved match to the data are associated with a larger value of ℒ (*θ* | *y*^obs^). It is convenient to work with the logarithm of the likelihood to avoid working with the product of many extremely small quantities, giving the log-likelihood function

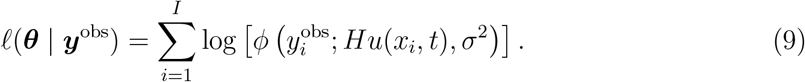

Note that maximising Equation (9) is mathematically identical to minimising the sum of squared differences (or equivalently the mean squared error) for constant *σ*^2^[48]. This equivalence means that very standard approaches of parameter estimation by minimising a least-squares objective function are equivalent to maximising the log-likelihood function under the assumption of working with an additive Gaussian noise model with constant variance.

Following the approach of Hines et al. [50, 51] we will demonstrate how to take a Bayesian approach to sampling this likelihood function which implicitly relies on Bayes’ theorem that we write as

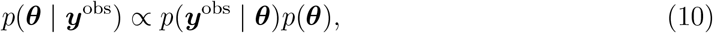

where *p*(*y*^obs^ | *θ*) is the likelihood function, *p*(*θ*) is the prior and *p*(*θ* | *y*^obs^) is the posterior distribution. Our aim is to estimate the posterior distribution, as it summarises information about the parameter *θ* in light of the data *y*^obs^ and the prior *p*(*θ*). The likelihood function describes information contributed by the data, corresponding to the probabilistic noise model, Equation (7), evaluated at the observed data. We explore the parameter space by sampling the posterior distribution using a Metropolis-Hastings Markov Chain Monte Carlo (MCMC) algorithm [50]. The Markov chain starts at position *θ*^(0)^ and a potential move to *θ*^*^is accepted with probability

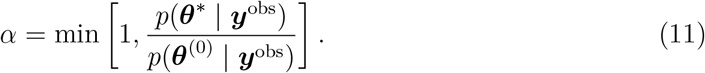

The Markov chain tends to move in the direction of high posterior probability, but also allows movements away from local minima by allowing occasional steps to regions of lower posterior probability [50, 51]. Markov chains produced by this algorithm explore the parameter space in proportion to the posterior probability. Once the MCMC chains have mixed, the sampler has converged to the target posterior distribution and is effectively exploring the full support of that distribution. Under these conditions the MCMC algorithm can be used to generate a finite number of approximately independent, identically distributed samples from the posterior distribution [50, 51]. Proposals in the MCMC algorithm are made by perturbing the current sample point by a draw from the multivariate normal distribution with zero mean and covariance matrix Σ. In this example with *θ* = (*U, D, σ*)^⊤^ we specify Σ = diag (10^−4^, 10^−4^, 10^−3^)^⊤^. The entries in the covariance matrix are hyperparameters for the MCMC algorithm, and the choice of these hyperparameters is critical because they can strongly affect the rate of mixing and convergence of the algorithm. In this case with a relatively small number of unknown parameters, we have selected these hyperparameters based on heuristic experience obtained using a number of pilot simulations [50, 51]. For higher dimensional problems it is possible to use adaptive methods to help in the selection of these hyperparameters [54, 55].

With this framework, we generate a Markov chain with 10^4^samples, starting at *θ*^(0)^ = (*U, D, σ*)^⊤^ = (0.1, 0.1, 0.1)^⊤^. The MCMC traces in Figure 3 show that samples of *θ* rapidly move away from the starting position and appear to settle into a stationary distribution after approximately 500-1000 samples. In this example we have chosen to stop the chain after 10^4^ samples as there is no obvious visually detectable drift at this point [50, 51]. The decision regarding when to stop the MCMC chains can be formalised by calculating certain criteria that involve comparing within-chain and between-chain variances [47]. For the purposes of this review we take a standard approach and stop the chain according to a simple visual inspection criteria [50, 51]. For typical applications of MCMC-based parameter estimation, we never know in advance of performing the potentially expensive MCMC simulation whether the chains will eventually mix and converge. If the chains do mix and converge, we never know in advance how many iterations will be required.

**Figure 3:**
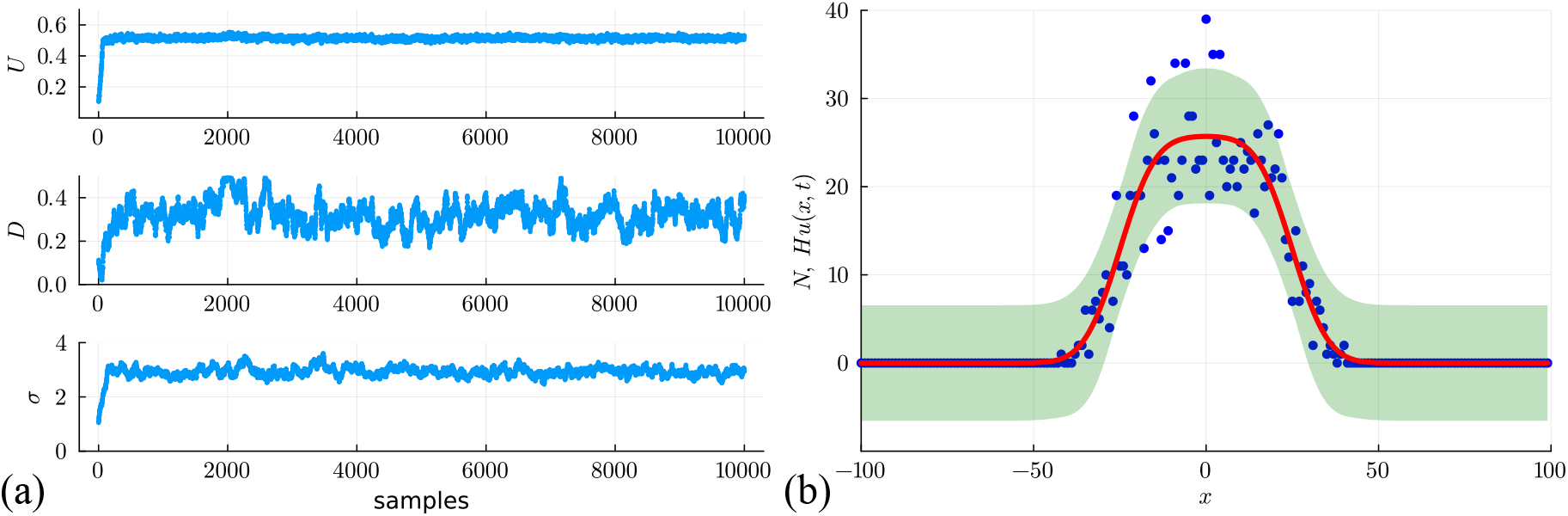
(a) MCMC trace plots for (*U, D, σ*)^⊤^ starting at *θ*^(0)^ = (*U, D, σ*)^⊤^ = (0.1, 0.1, 0.1)^⊤^. Proposals are drawn from the multivariate normal distribution with zero mean and covariance matrix Σ = diag (10^−4^, 10^−4^, 10^−3^)^⊤^. The final 2000 samples are taken as samples from the stationary distribution, and the mean of these samples gives point estimates (*U, D, σ*)^⊤^ = (0.515, 0.332, 2.959)^⊤^. (b) Posterior prediction interval generated using the final *M* = 1000 samples from (a) (green shaded) encloses 94.0% of the stochastic data. This interval is superimposed on the data (blue dots) and the solution of the continuum model *Hu*(*x, t*), evaluated at the posterior mean (*U, D, σ*)^⊤^ = (0.515, 0.332, 2.959)^⊤^ (solid red).

To quantify the results of the MCMC chain we take the final 2000 samples from chain and compute various marginal distributions summarised in Figure 4 where we see that each of the univariate posterior marginal distributions are unimodal and relatively narrow about the peak. Accordingly we characterise these distributions by their mean, which gives point estimates of (*U, D, σ*)^⊤^ = (0.515, 0.332, 2.959)^⊤^, confirming that our estimates of *U* and *D* are relatively close to the expected result, as well as being consistent with the point estimates and credible intervals obtained using ABC rejection. The MCMC-derived credible intervals are relatively narrow, indicating that our estimates are reasonably precise. Figure 4 also presents histograms illustrating the bivariate marginal distributions between each pair of parameters which gives us a sense of any correlation between parameter pairs. The shapes of the distributions indicate no obvious correlation between *D* and *σ* or between *U* and *σ*. In contrast there appears to be some degree of positive correlation between *D* and *U*, which is consistent with the previous ABC rejection results in Figure 2(b).

**Figure 4:**
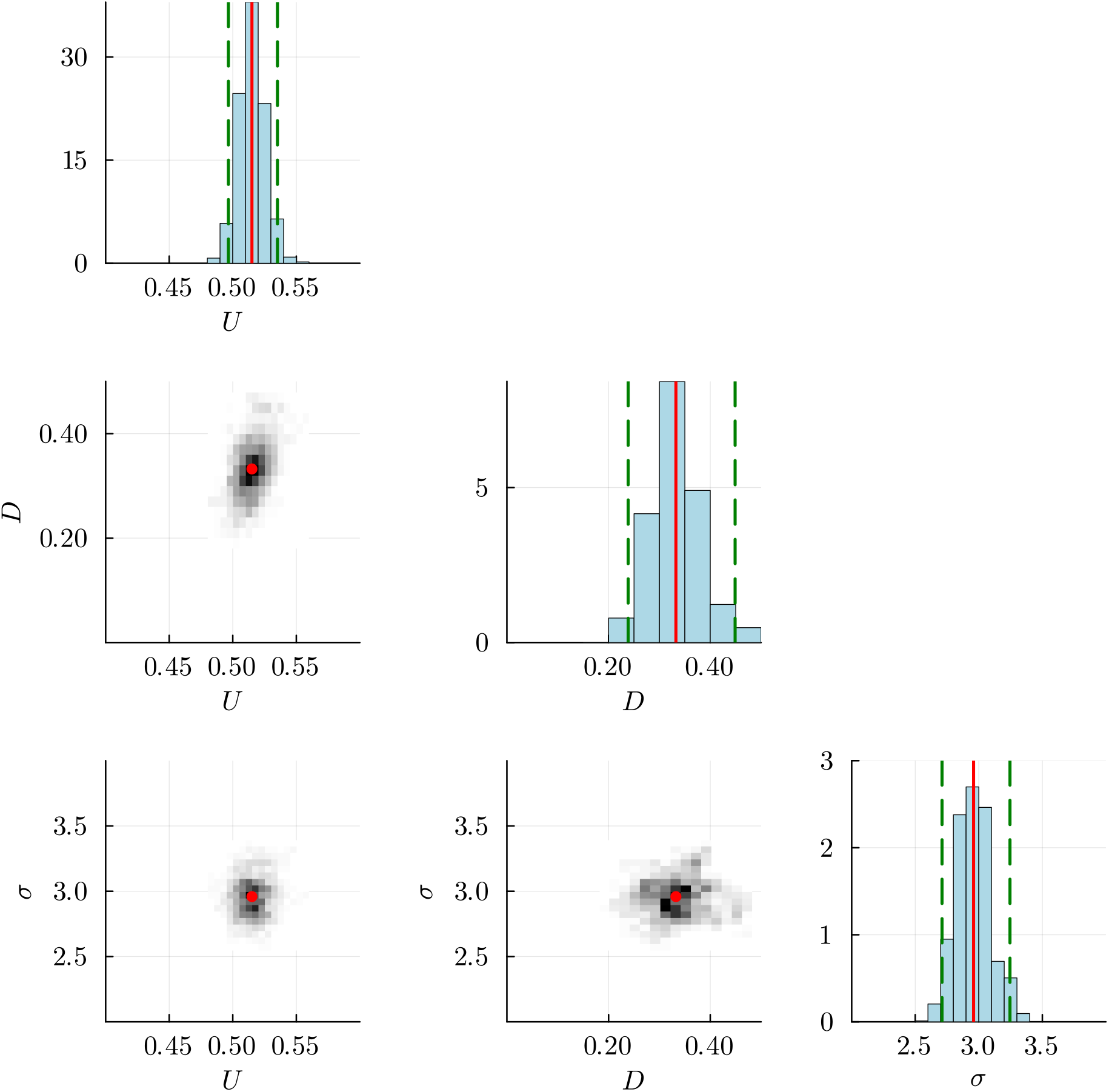
Plot matrix summary constructed from the final 2000 MCMC samples in Figure 3(a). Diagonal entries show histograms of marginal posterior distribution, where each histogram is superimposed with the mean (red vertical lines) and the 0.025 and 0.975 quantiles of the distributions (green dashed vertical lines). These correspond to estimates of *U* = 0.515 [0.497, 0.535]; *D* = 0.332 [0.238, 0.449]; and, *σ* = 2.959 [2.711, 3.245]. Off diagonal plots show histograms representing the joint posterior distributions between pairs of parameters with the point estimates superimposed (red dot).

Since our MCMC chain seems to have converged to produce relatively narrow posterior distributions, we can consider generating a posterior predictive interval [9, 10] with the additional information provided through incorporating the additive Gaussian noise model and our estimates of *σ*. To proceed we take the final *M* = 1000 samples from the Markov chain in Figure 3, and for each set of parameter values we solve the PDE model, giving *M* solutions denoted *Hu*_*m*_(*x, t*) for *m* = 1, 2, 3, …, *M*. The quantity *Hu*(*x, t*) represents the mean of the measurement error model at position *x* and time *t*. With the measurement error model, we no longer need to restrict ourselves to working with the mean of that distribution as we can now incorporate information about the width of the noise model given the estimates of *σ* from the Markov chain. Therefore, at the *i*th spatial location, we consider an interval [*Hu*_*m*_(*x, t*)^−^, *Hu*_*m*_(*x, t*)^+^], where the lower and upper values correspond to the 0.025 and 0.975 quantiles of the estimated noise distribution, respectively. We construct the posterior predictive interval, as before, except that instead of taking the union of point estimates at each point *x*_*i*_, we now take the union of the intervals [*Hu*_*m*_(*x*_*i*_, *t*)^−^, *Hu*_*m*_(*x*_*i*_, *t*)^+^] over *m* = 1, …, *M* at each *x*_*i*_. The resulting prediction interval is given in Figure 3(b) where we also superimpose the data and the solution of the model evaluated at the posterior mean (*U, D, σ*)^⊤^ = (0.515, 0.332, 2.959)^⊤^. The prediction interval takes into account the uncertainty in the model parameters *θ* = (*U, D, σ*)^⊤^ as well as the width of the distribution associated with the measurement error model. The solution of the model evaluated at the posterior mean captures key trends in the noisy data, and the prediction interval captures much of the variability of the data. One drawback of working with the additive Gaussian noise model is that the measurement error model has a constant variance, and so the width of the prediction interval is approximately constant for all *x*. This means that in regions where *u*(*x, t*) is sufficiently close to zero the prediction interval leads to a non-physical negative lower bound, *Hu*_*m*_(*x*_*i*_, *t*)^−^ *<* 0. The fact that the measurement noise model is mis-specified could also bias estimates of the process model parameters *D* and *U* [39, 53]. We will describe ways of addressing this issue in Section 3.3.

### 3.3 Likelihood-based optimisation: Estimation and prediction

We now explore a fourth and final approach to estimate *θ* = (*U, D, σ*)^⊤^ from the noisy count data in Figure 1(f). By continuing to work with the additive Gaussian noise model, we have access to the loglikelihood function given by Equation (9) [48]. Unlike the approaches in Sections 3.1–3.2 that rely either on repeated simulations of the stochastic model, or repeated evaluation of the solution of the surrogate model via MCMC sampling, here we take a more direct approach by numerically maximising *ℓ*(*θ* | *y*^obs^) to provide point estimates of *θ* = (*U, D, σ*)^⊤^. Using a similar approach based on numerical optimisation, we then construct a series of univariate profile loglikelihood functions [48, 56]. This provides a rapid means of assessing the degree of curvature of the loglikelihood function, which provides key information about inferential precision and parameter identifiability, as we now discuss.

Identifiability is a property which a model must satisfy for precise parameter inference, and identifiability analysis refers to a group of methods that determine how well the model parameters can be estimated given the observed data [48, 57]. Methods of identifiability analysis are typically classified in terms of dealing with *structural* or *practical* identifiability [50]. Structural identifiability analysis determines whether different parameter values generate different probability distributions of the observable variables [58, 59], with the implication that given access to an infinite amount of perfect, noise-free data, it would then be possible to precisely estimate the model parameters. Alternatively, practical identifiability analysis involves working with noisy, sparse data, and exploring the extent to which parameter values can be confidently estimated, given these data. Practical identifiability analysis usually involves fitting a mathematical model to data, and then exploring the extent to which the model fit changes as the parameters vary [48]. A common tool for assessing practical identifiability is the profile likelihood [23, 24, 48, 60].

Instead of repeated sampling of the posterior distribution, our fourth approach to inference and prediction involves applying maximum likelihood estimation (MLE) to obtain the best-fit set of parameters,

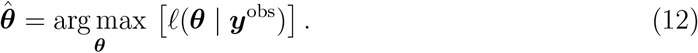

All numerical optimisation computations in this review use the Nelder-Mead algorithm with simple bound constraints [61, 62], and all computations are performed using default stopping criteria within the well-known NLopt family of numerical optimisation routines [63]. Given 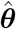, we then work with the normalised loglikelihood function 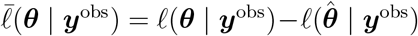, so that 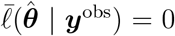. Working with 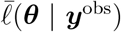 allows us to define asymptotic confidence sets by using Wilks’ theorem [64], which states that the likelihood ratio statistic converges in distribution to a *χ*^2^ variable [8, 56, 64]. This result enables us to construct confidence regions and confidence sets, thereby allowing us to quantitatively explore how a noisy set of data can be related to variability in parameter estimates. A key advantage of working with Wilks’ theorem is that this theorem holds for arbitrary dimensions and parameter reparameterisations. A limitation of working with Wilks’ theorem is that it is an asymptotic result that may not hold when dealing with small sample sizes [8]. In practice, Wilks’ theorem allows us to define a likelihood-based threshold, 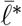, that defines confidence regions in parameter space 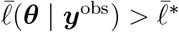, and corresponding confidence sets, 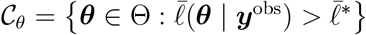. We will explain how to compute values of 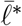 later. [8, 56].

To assess parameter identifiability we take a profile likelihood-based approach [22, 56, 60, 65– 68] by partitioning *θ* into interest parameters *ψ*, and nuisance parameters *ω*, so that *θ* = (*ψ, ω*)^T^. For simplicity, here we always take the interest parameter to be a single parameter which allows us to construct various univariate profile likelihood functions. Regardless, for a set of data *y*^obs^, the profile loglikelihood for the interest parameter *ψ* given the partition (*ψ, ω*)^T^is

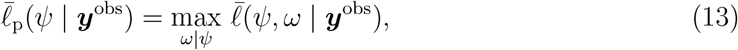

which implicitly defines a function *ω*^*^(*ψ*) of optimal values of *ω* for each value of *ψ*, and defines a surface with points (*ψ, ω*^*^(*ψ*)) in parameter space. The targeting property of the profile likelihood allows us to explore the identifiability of each parameter which is particularly insightful and can be computationally efficient, especially compared to MCMC-based approaches, which normally characterise non-identifiability by poorly mixing MCMC chains [69, 70]. For univariate profiles, 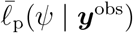 is a function of one parameter that can be plotted and visualised in the usual way. For example, considering the current problem with *θ* = (*U, D, σ*)^⊤^, we can construct a univariate profile likelihood function for *U* by setting *ψ* = *U* and *ω* = (*D, σ*)^⊤^. As for calculating the MLE, profile likelihoods are computed using the Nelder-Mead algorithm to evaluate the necessary numerical optimisation problems [63], and in all cases the iterative optimisation calculations can be initialised using the appropriate components of 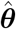, and all profile likelihood functions are computed across an interval of the interest parameter that is uniformly discretised.

For the data in Figure 1(e) and the loglikelihood function given by Equation (9), numerical optimisation gives 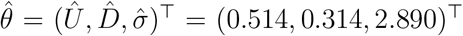 within a fraction of a second of computation time. These estimates compare very well those obtained by repeated sampling via the MCMC algorithm in Figure 3. In contrast, however, the optimisation-based estimates can be obtained with several orders of magnitude computational speed-up, and are obtained without the need for specifying hyperparameters for the MCMC parameter proposals. To assess the identifiability of these point estimates, the univariate profile log-likelihood functions for *U, D*, and *σ* are given in Figure 5(a). We see that each profile is characterised by a unique maximum at the MLE. The profiles are sufficiently curved that we can construct an approximate confidence interval for each parameter by finding the value of the parameter where the profile loglikelihood intersects with the 95% asymptotic threshold where 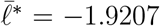 as we explain further below. For example, in terms of the initial density we have *Û*= 0.514, with an approximate 95% confidence interval of *U* ∈ [0.494, 0.535]. Just like the MLE calculation, the construction of univariate profile likelihood functions is computationally efficient relative to MCMC sampling [50, 51, 70]. In general, we never know in advance how curved the univariate profile likelihood function will be before we calculate that function. Depending on the degree of curvature of the profile likelihood function, it is possible to conclude that a parameter is practically identifiable, poorly-identified, or one sided identifiable [71]. This judgement is usually made on a case-by-case basis depending on the width of the approximate confidence interval relative to problem-specific constraints and expectations [72].

**Figure 5:**
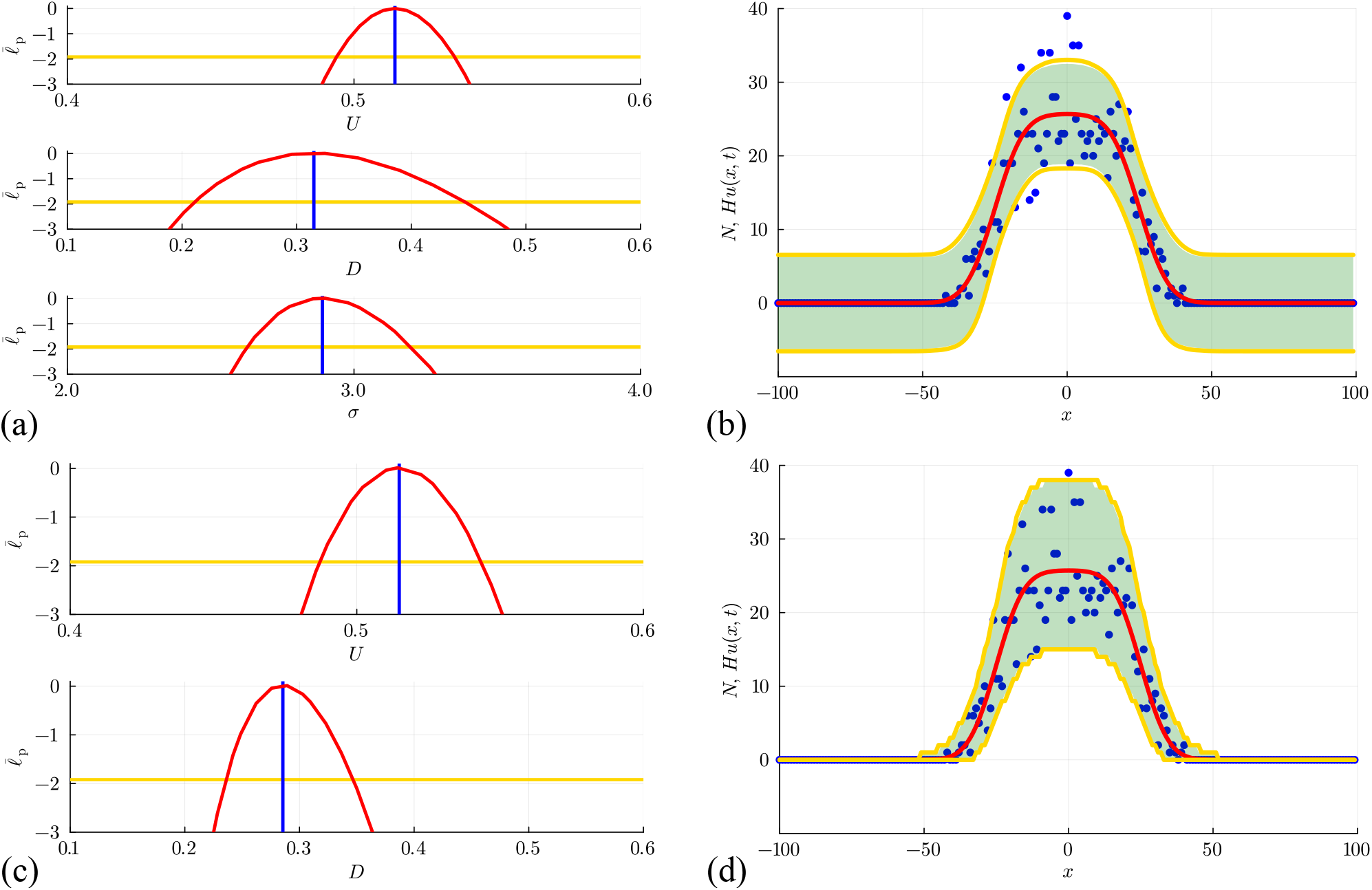
(a)–(b) Estimation, identifiability analysis and prediction under the additive Gaus _sian noise model. (a) Profiles for *U, D* and *σ* with 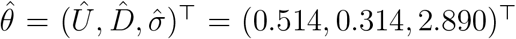_ superimposed (vertical blue lines). Each univariate profile gives an approximate 95% con _fidence interval: *Û* = 0.514 [0.494, 0.535]; 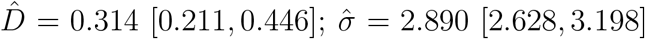_ generated by finding the values of the interest parameter where the profile intersects with the 95% threshold (yellow horizontal lines). (b) Prediction interval generated with the additive Gaussian noise model encloses 95.0% of the data. (a)–(b) Estimation, identifiabil ity analysis and prediction under the Poisson noise model. (c) Profiles for *U* and *D* with 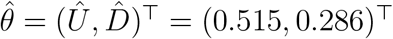. Each univariate profile gives an approximate 95% confidence _interval: *Û* = 0.515 [0.487, 0.543]; 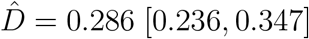_. (d) Prediction interval generated with the Poisson noise model encloses 99.5% of the data. The prediction intervals in (b) and (d) compare the approach of uniform sampling (solid green region) with the interval generated using Laplace’s approximation (solid yellow bounds).

Once we have established the normalised loglikelihood function, we define asymptotic confidence sets by identifying a likelihood-based threshold,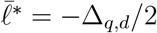,where Δ_*q,d*_ refers to the *q*th quantile of the *χ*^2^ distribution with *d* degrees of freedom [8, 64]. The asymptotic confidence set is the set of values of *θ* satisfying 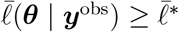. In this work we use a 0.95 quantile threshold for problems with *d* = 1, 2, 3 and 4 degrees of freedom, for which the threshold values are 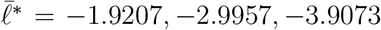 and −4.7439, respectively. In this context, the number of degrees of freedom corresponds to the number of free parameters [73]. Given 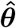 and confirmation via the profile likelihood function that these estimates are well-identified by the data, we now compute a prediction interval based on these data. We will construct a prediction interval [74] using two methods. First, we generate *M* = 1000 samples of *θ* that lie within the approximate 95% confidence region where 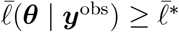. These samples are generated using a simple rejection algorithm where parameters are uniformly randomly sampled within a user-defined region containing 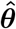. With these *M* parameter samples we can solve the mathematical model to give *M* continuous solution curves *Hu*_*m*_(*x, t*) for *m* = 1, 2, 3 …, *M*. At each spatial location *x* we compute an interval [*Hu*_*m*_(*x, t*)^−^, *Hu*_*m*_(*x, t*)^+^], where the lower and upper values correspond to the 0.025 and 0.975 quantiles of the noise model with mean *Hu*(*x, t*), respectively. As before, we take the union of intervals [*Hu*_*m*_(*x*_*i*_, *t*)^−^, *Hu*_*m*_(*x*_*i*_, *t*)^+^] over *m* = 1, 2, 3, …, *M* at each spatial location *x*_*i*_ on a uniform discretisation of the domain, −100 ≤ *x* ≤ 100 using 201 mesh points to give the green shaded prediction interval in Figure 5(b). As in Section 3.1, we see that the solution evaluated at the MLE captures the main trends and features of the count data, but now we see the value in introducing the noise model as here the prediction interval encloses the stochastic data reasonably well.

Our approach of using uniformly random samples to find *M* parameter values lying within the region where 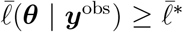 is typically the most computationally-expensive aspect of the workflow in Figure 5(a)–(b). The computational overhead of this step depends upon the shape of 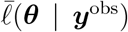, the number of free parameters, and the user-defined size of the region about 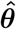 from which random samples are drawn. In our experience, working with this approach requires user-derived expertise in determining the extent of the region about 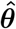 from which random samples are drawn. Choosing a region that is too narrow means that the samples do not reach the asymptotic limit given by Wilks’ theorem, whereas choosing a region that is too wide leads to very slow sampling.

Another approach involves invoking Laplace’s approximation where the shape of the loglikelihood function is approximated with a multivariate normal distribution [9, 10, 75–77]. To implement this approximation we compute the observed Fisher Information at the MLE,

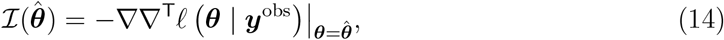

which is a square Hessian matrix, whose elements are the second order partial derivatives of *ℓ* with respect to the elements of *θ*. In this study we compute the elements of 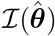 using automatic differentiation [78, 79], which avoids working with finite difference approximations and truncation error [80]. Given our estimates of 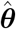 and 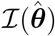, we approximate the shape of the loglikelihood function in the vicinity of the MLE as a multivariate normal distribution, 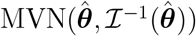. With this approximation we can draw *M* = 1000 random samples of 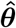 from this multivariate normal distribution. As before, we then calculate the PDE solution *u*_*m*_(*x, t*) and construct an interval [*Hu*_*m*_(*x, t*)^−^, *Hu*_*m*_(*x, t*)^+^] for each sample *m* = 1, …, *M*. We then define the prediction interval to be from the 0.025 quantile of *Hu*_*m*_(*x, t*)^−^ to the 0.975 quantile of *Hu*_*m*_(*x, t*)^+^.

It is important to note that drawing samples from a uniform distribution in the region where 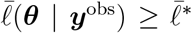 is different to drawing samples from the multivariate normal distribution given by Laplace’s approximation, which has more density close the MLE and some density in the tails of the distribution. In practice we find that the computational effort is significantly lower for the second approach. These two methods give prediction intervals that are very similar, as shown in Figure 5(b). This comparison illustrates that working with Laplace’s approximation can give accurate prediction interval estimation with a significantly reduced computational overhead. It should be noted that the intervals defined by both of these methods are technically known as *95%/95% tolerance intervals*, meaning that they are expected to cover 95% of the observation distribution in at least 95% of independent repeats of the same experiment. This is somewhat wider than a 95% prediction interval, which is expected to contain a single future observation with 95% probability [74]. However, in keeping with much of the literature in this area, we will refer to these as prediction intervals throughout this review.

Before we consider moving on to explore different types of data generated by different random walk models, we briefly re-examine the data in Figure 1(f) using a likelihood-based optimisation approach. In this instance, however, we work with a different noise model to address some of the challenges associated with the commonly-used additive Gaussian noise model underlying results in Figure 5(a)–(b). Given that our data consists of non-negative integer count data, and we express the solution of our continuum limit model as *Hu*(*x, t*), which we interpret as the expected number of agents at location *x* and time *t*, an alternative and perhaps more natural interpretation of this situation is that the data are drawn from a Poisson distribution whose mean is *Hu*(*x, t*), where *u*(*x, t*) is the solution of the continuum limit model

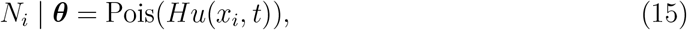

at all locations *x*_*i*_for *i* = 1, 2, 3, …, *I*. As we will now explore, there are several advantages, both conceptual and procedural, in working under this Poisson-based noise model. First, by definition, the Poisson distribution has support on non-negative integers which is well-suited to the non-negative integer count data generated by the random walk model or a biological experiment. Second, the Poisson noise model does not involve specifying another parameter which means that here we estimate *θ* = (*U, D*)^⊤^ instead of *θ* = (*U, D, σ*)^⊤^ like we did with the additive Gaussian noise model. With this framework numerical optimisation gives 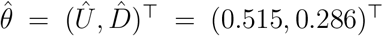, which compares well with the estimates obtained using the additive Gaussian noise model either by sampling or optimisation. To assess the identifiability of these point estimates, the univariate profile loglikelihood functions for *U* and *D* are given in Figure 5(c), where we see that each parameter is well-identified by the data as the profiles are characterised by a unique maximum at the MLE, and are sufficiently curved that we can construct an approximate confidence interval for each parameter by estimating the intersection of the univariate profile log-likelihood and the 95% asymptotic threshold [73]. These results are much faster to obtain than the MCMC sampling-based results in Figure 3, as well as being modestly faster to obtain than the optimisation results in Figure 5(a)–(b) for the additive Gaussian noise model since we are dealing with a reduced number of unknown parameters.

Perhaps the most obvious advantage in using the Poisson noise model is clear in terms of the prediction intervals summarised in Figure 5(d). As before, we obtain the prediction intervals using the same two methods used in Figure 5(b). Both methods give similar results to each other, but working with Laplace’s approximation leads to an important computational speed-up. Importantly, regardless of how we sample *θ*, the Poisson-based prediction intervals are very different from those in Figure 5(b). In the former case we have a prediction interval with constant width that gives the misleading impression that negative count data are possible in regions where *u*(*x, t*) is sufficiently close to zero. In contrast, the Poisson-based prediction intervals completely avoids the possibility of negative counts, as the width of the Poisson-based prediction interval increases with *u*(*x, t*), and vanishes as *u*(*x, t*) → 0^+^. This feature gives a much more realistic prediction interval that reflects the fluctuations in the data much more accurately. In this case, the prediction intervals obtained using the additive Gaussian noise model encloses 95.0% of the data, and fails to enclose several data points where the expected counts are high in the centre of the lattice, whereas the Poisson-based prediction interval encloses 99.5% of the data.

In summary, working with the Poisson noise model increases the computational efficiency with which we can estimate parameters, assess their identifiability, and perform model predictions. This approach also increases model fidelity since it leads to predictions that are more realistic and appropriate for count data than making the standard assumption that the data are distributed according to a normal distribution with constant variance. While it is also possible to use the Poisson noise model to repeat the exercise in Figure 3 and use MCMC sampling, we will now move on to a set of new problems by exploring a range of combinations of different random walk models and measurement error models. For the remainder of this review we will focus on the most computationally efficient approaches, namely likelihood-based optimisation approaches for parameter estimation and identifiability analysis, along with invoking Laplace’s approximation using automatic differentiation to generate parameter samples and construct prediction intervals.

### 3.4 Combining different process and noise models

To explore the generality of the approaches considered so far, we will now extend the process model and explore our ability to estimate parameters, understand their identifiability, and make model-based predictions for a range of different models. To keep track of the various combinations of process and noise models we summarise the cases in Table 1. To begin we extend the random walk model used in Sections 3.1–3.3 to now include both biased motion [20, 81] and agent proliferation [21]. The random walk model is implemented on the same lattice with the same sequential update method. To evolve the system from time *t* to time *t* + *τ*, we consider a random sequential update method where for a population of *Q* agents, we select *Q* agents independently at random, one at a time, with replacement. Each selected agent is given the opportunity to move with probability *P* ∈ [0, 1]. If an agent at site (*i, j*) is selected to move, the target site is chosen with a directional bias. Lattice sites (*i, j ±* 1) are chosen with probability 1*/*4, whereas sites (*i ±* 1, *j*) are chosen with probability (1 *± ρ*)*/*4. Here *ρ* ∈ [−1, 1] is a constant bias parameter such that setting *ρ* = 0 gives an unbiased random walk, setting *ρ >* 0 means that agents are more likely to step in the positive *x*-direction, and setting *ρ <* 0 means that agents are more likely to step in the negative *x*-direction. Once *Q* potential motility events have been considered, we then select *Q* agents independently at random, one-at-a-time, with replacement. Each selected agent proliferates with probability *R* ∈ [0, 1]. If an agent at site (*i, j*) proliferates we simply place another agent at the same site. Once *Q* motility events and *Q* proliferation events have been attempted we update our value of *Q* and the current time step is completed.

**Table 1:** Summary of 12 inference and prediction cases considered in this review.

| Case | Exclusion | Bias | Growth | Flux | Source | Noise |
| --- | --- | --- | --- | --- | --- | --- |
| 1: Figure 5(a)-(b) | No | No | No | $J = -D \frac{\partial u}{\partial x}$ | $S = 0$ | Additive Gaussian |
| 2: Figure 5(c)-(d) | No | No | No | $J = -D \frac{\partial u}{\partial x}$ | $S = 0$ | Poisson |
| 3: Figure 7(a)-(b) | No | Yes | No | $J = -D \frac{\partial u}{\partial x} + vu$ | $S = 0$ | Additive Gaussian |
| 4: Figure 7(c)-(d) | No | Yes | No | $J = -D \frac{\partial u}{\partial x} + vu$ | $S = 0$ | Poisson |
| 5: Figure 9(a)-(b) | No | No | Yes | $J = -D \frac{\partial u}{\partial x}$ | $S = ru$ | Additive Gaussian |
| 6: Figure 9(c)-(d) | No | No | Yes | $J = -D \frac{\partial u}{\partial x}$ | $S = ru$ | Poisson |
| 7: Figure 11(a)-(b) | Yes | No | No | $J = -D \frac{\partial u}{\partial x}$ | $S = 0$ | Additive Gaussian |
| 8: Figure 11(c)-(d) | Yes | No | No | $J = -D \frac{\partial u}{\partial x}$ | $S = 0$ | binomial |
| 9: Figure 12(a)-(b) | Yes | Yes | No | $J = -D \frac{\partial u}{\partial x} + vu(1 - u)$ | $S = 0$ | Additive Gaussian |
| 10: Figure 12(c)-(d) | Yes | Yes | No | $J = -D \frac{\partial u}{\partial x} + vu(1 - u)$ | $S = 0$ | binomial |
| 11: Figure 13(a)-(b) | Yes | No | Yes | $J = -D \frac{\partial u}{\partial x}$ | $S = ru(1 - u)$ | Additive Gaussian |
| 12: Figure 13(c)-(d) | Yes | No | Yes | $J = -D \frac{\partial u}{\partial x}$ | $S = ru(1 - u)$ | binomial |

The mean behaviour of the discrete model is described by a continuous linear PDE that can be written as [20, 21, 31, 32],

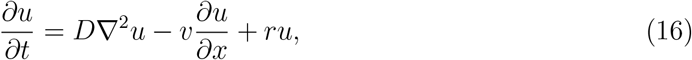

where 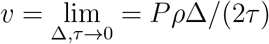 and 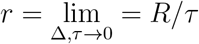, where *u*(*x, y, t*) ≥ 0 denotes the average density of agents at location (*x, y*) at time *t*. Again, since our simulations do not involve macroscopic density gradients in the vertical direction the PDE simplifies to

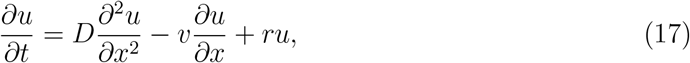

for the density *u*(*x, t*) ≥ 0. With our choice of initial conditions, and simulations over an intermediate time scale where agents do not touch the vertical boundaries at *x* = *± L*, we can approximate the solution of Equation (17) with the appropriate infinite domain solution [40]

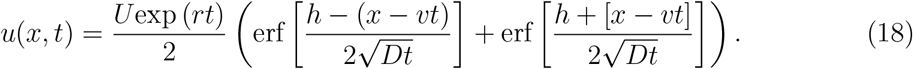

To illustrate we present a simulation in Figure 6 where we have the same initial condition as before, but the population evolves with *P* = 1, *ρ* = 0.5 and *R* = 0, which is associated with *D* = *v* = 0.25 and *r* = 0. The drift in the positive *x*-direction is visually obvious in the distribution of agents in Figure 6(b) as well as being reflected in the count data in Figure 6(d). As for the unbiased simulation in Figure 1, the solution of the mean-field PDE model, Equation (18), provides a good match to the mean count data, but does not capture the stochastic fluctuations generated by the stochastic model.

**Figure 6:**
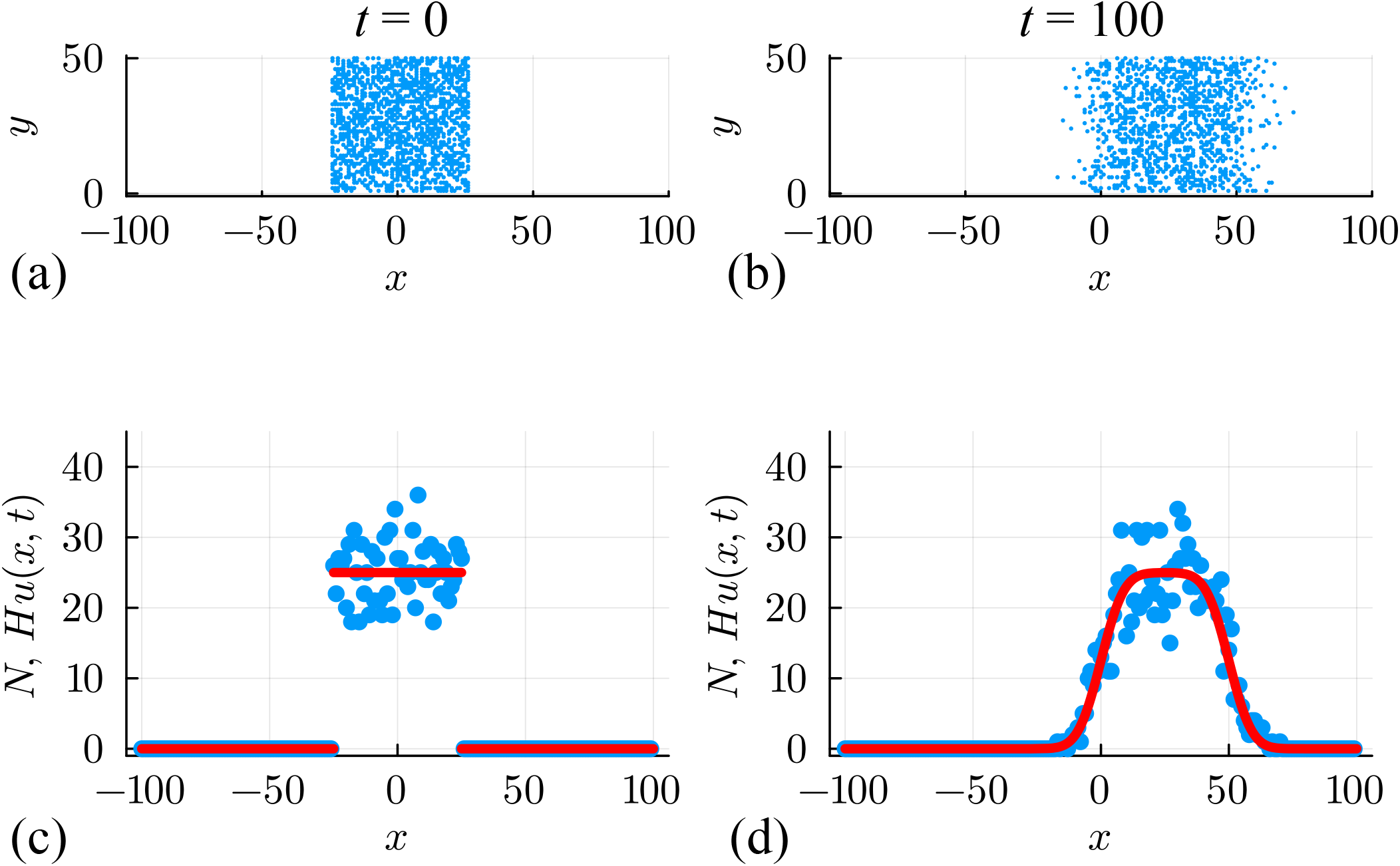
(a)–(b) Biased non-interacting random walk simulation on a square lattice with Δ = *τ* = 1, *H* = 50 and *L* = 100. The initial placement of agents is shown in (a), and the outward spreading of agents with a drift in the positive *x*-direction after 100 time steps is shown in (b). The initial placement of agents is the same as in Figure 1, and random agent migration takes place with *P* = 1 and *ρ* = 0.5. (c)–(d) Numbers of agents per column at *t* = 0 and *t* = 100, respectively. Noisy count data (blue dots) is superimposed with *Hu*(*x, t*), where *u*(*x, t*) is the solution of Equation (17) with *D* = *P/*4, *v* = *P ρ/*2 and *r* = 0 (red curve).

We first attempt to use data in Figure 6 to estimate parameters, examine parameter identifiability, and construct prediction intervals by making the standard assumption that the count data is normally distributed about *Hu*(*x, t*) with constant variance. Numerical optimisation gives 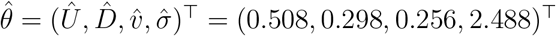, where the estimates of *U, D* and *v* that are close to the expected values. Four univariate profile likelihood functions in Figure 7(a) confirm that these parameters are well-identified by these data. The resulting prediction interval in Figure 7(b) encloses 95.0% of the noisy data, but again suffers from the limitation that the additive Gaussian noise model with constant variance leads to prediction intervals that include negative counts in regions where *u*(*x, t*) is sufficiently close to zero, and fails to enclose some of the data in regions where *u*(*x, t*) is sufficiently large.

**Figure 7:**
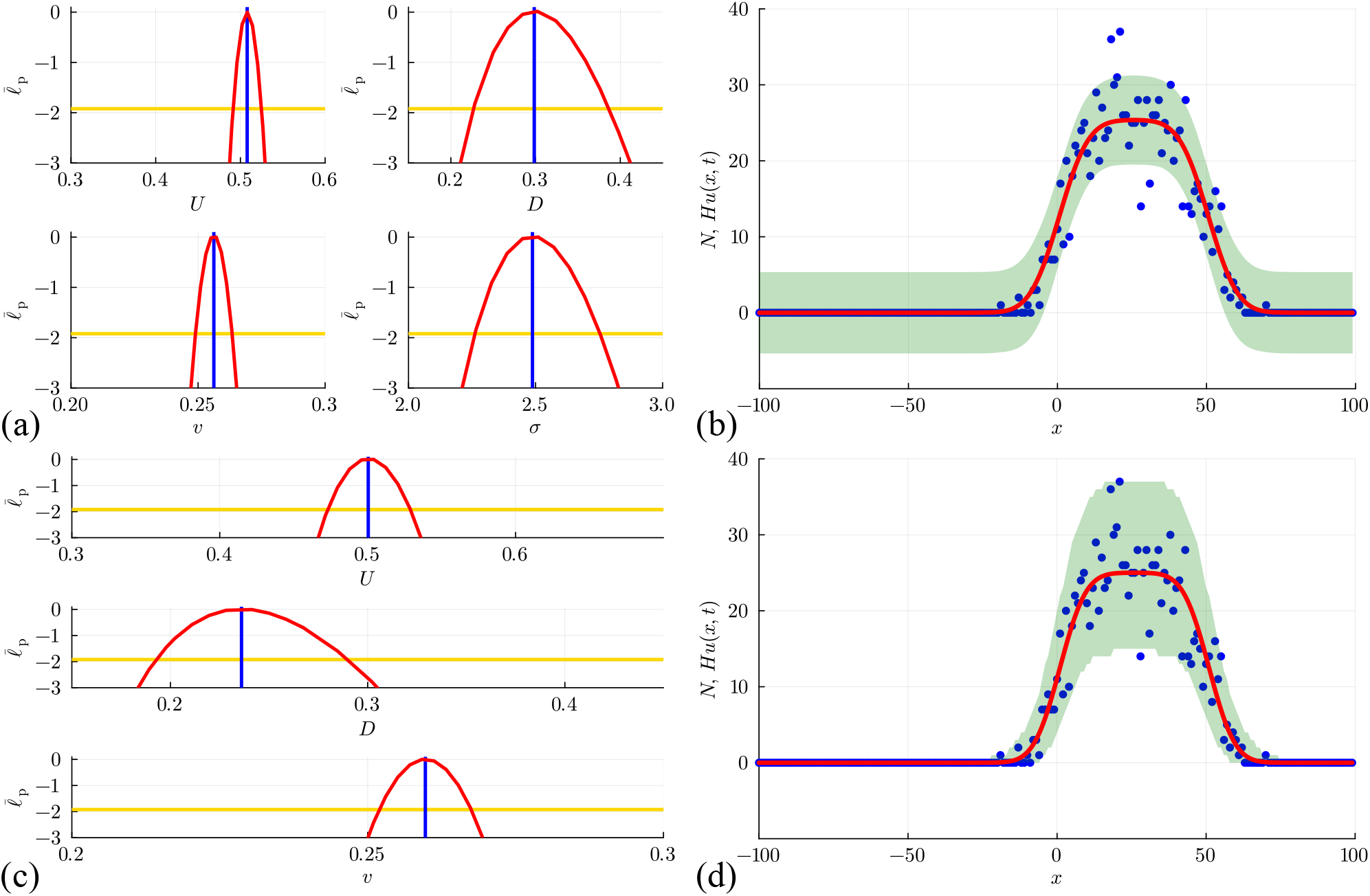
(a)–(b) Estimation, identifiability analysis and prediction under the additive _Gaussian noise model. (a) Profiles for *U, D, v* and *σ* with 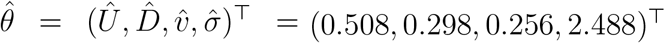_ superimposed (vertical blue lines). Each univariate profile gives _an approximate 95% confidence interval: *Û* = 0.508 [0.491, 0.525]; 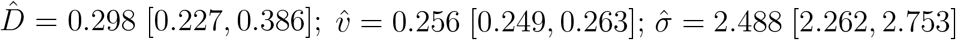_ generated by finding the values of the interest parameter where the profile intersects with the 95% threshold (yellow horizontal lines). (b) Prediction interval generated with the additive Gaussian noise model encloses 95.0% of the data. (c)–(d) Estimation, identifiability analysis and prediction under the Poisson _noise model. (c) Profiles for *U, D* and *v* with 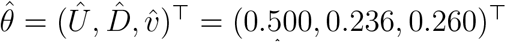_. Each _univariate profile gives an approximate 95% confidence interval: *Û*_ = 0.500 [0.473, 0.529]; 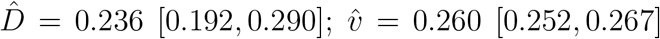. (d) Prediction interval generated with the Poisson noise model encloses 99.0% of the data. Prediction intervals in (b) and (d) are generated using Laplace’s approximation.

Further results in Figure 7(c)–(d) repeat the same estimation, identifiability, and prediction _steps under the Poisson noise model, leading to 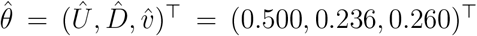_, which is close to the expected result, and each univariate profile likelihood function indicates that all three parameters are practically identifiable. The prediction interval in Figure 7(d) reflects the positive count data more accurately than the results obtained with the additive Gaussian noise model and encloses 99.0% of the data.

An additional simulation in Figure 8 considers the same initial condition as before, except that now agents undergo unbiased movement and proliferation with *P* = 1, *ρ* = 0, and *R* = 0.01, which is associated with *D* = 0.25 and *r* = 0.01. The population growth is clear in the count data in Figure 6(d), where we see that *Hu*(*x, t*) *> Hu*(*x*, 0) = 50 near *x* = 0 after 100 time steps. Estimates of *Hu*(*x, t*) obtained using Equation (18) provide a good match to the mean count data without capturing stochastic fluctuations. In this case with proliferation we will treat *U* as a known quantity and aim to estimate *D, r* and *σ*, and we will discuss this choice later in Section 6. We estimate parameters, test their identifiability and construct prediction intervals under the standard assumption that count data are normally distributed about *Hu*(*x, t*) with constant variance, giving 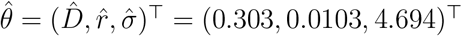. Three univariate profile likelihood functions in Figure 9(a) confirm that these parameters are well identified by this data. The prediction interval in Figure 9(b) encloses 93.5% of the data and suffers from the same limitation as before that the prediction intervals include negative counts in regions where *u*(*x, t*) is sufficiently close to zero, and fails to enclose some of the data in regions where *u*(*x, t*) is sufficiently large.

**Figure 8:**
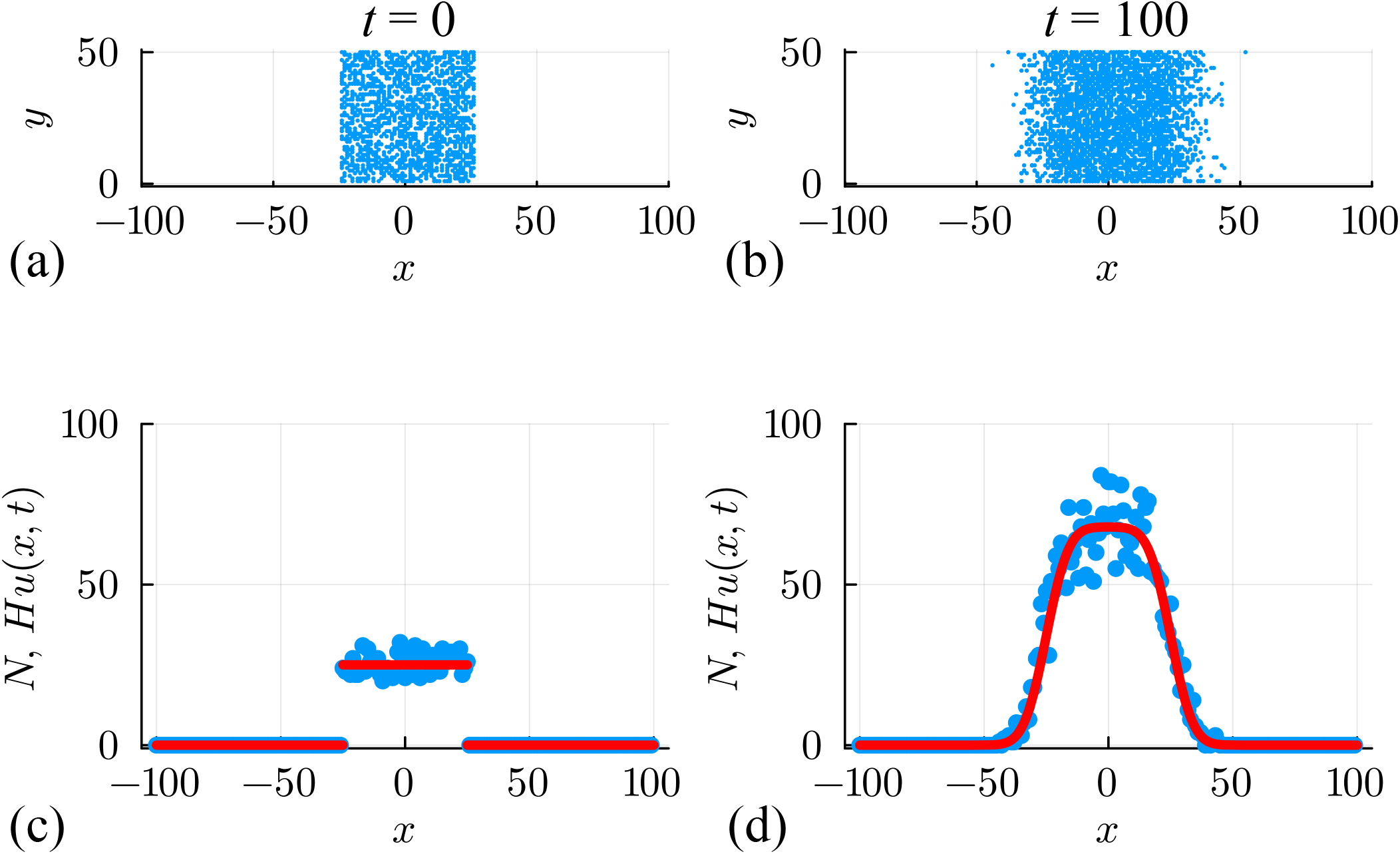
(a)–(b) Unbiased non-interacting random walk simulation with growth on a square lattice with Δ = *τ* = 1, *H* = 50 and *L* = 100. The initial placement of agents is shown in (a), and the combined spreading and growth of the population after 100 time steps is shown in (b). The initial placement of agents is the same as in Figure 1, and random agent migration and growth takes place with *P* = 1 and *R* = 0.01. (c)–(d) Numbers of agents per column at *t* = 0 and *t* = 100, respectively. Noisy count data (blue dots) is superimposed with *Hu*(*x, t*), where *u*(*x, t*) is the solution of Equation (17) with *D* = *P/*4, *v* = 0 and *r* = *R* (red curve).

**Figure 9:**
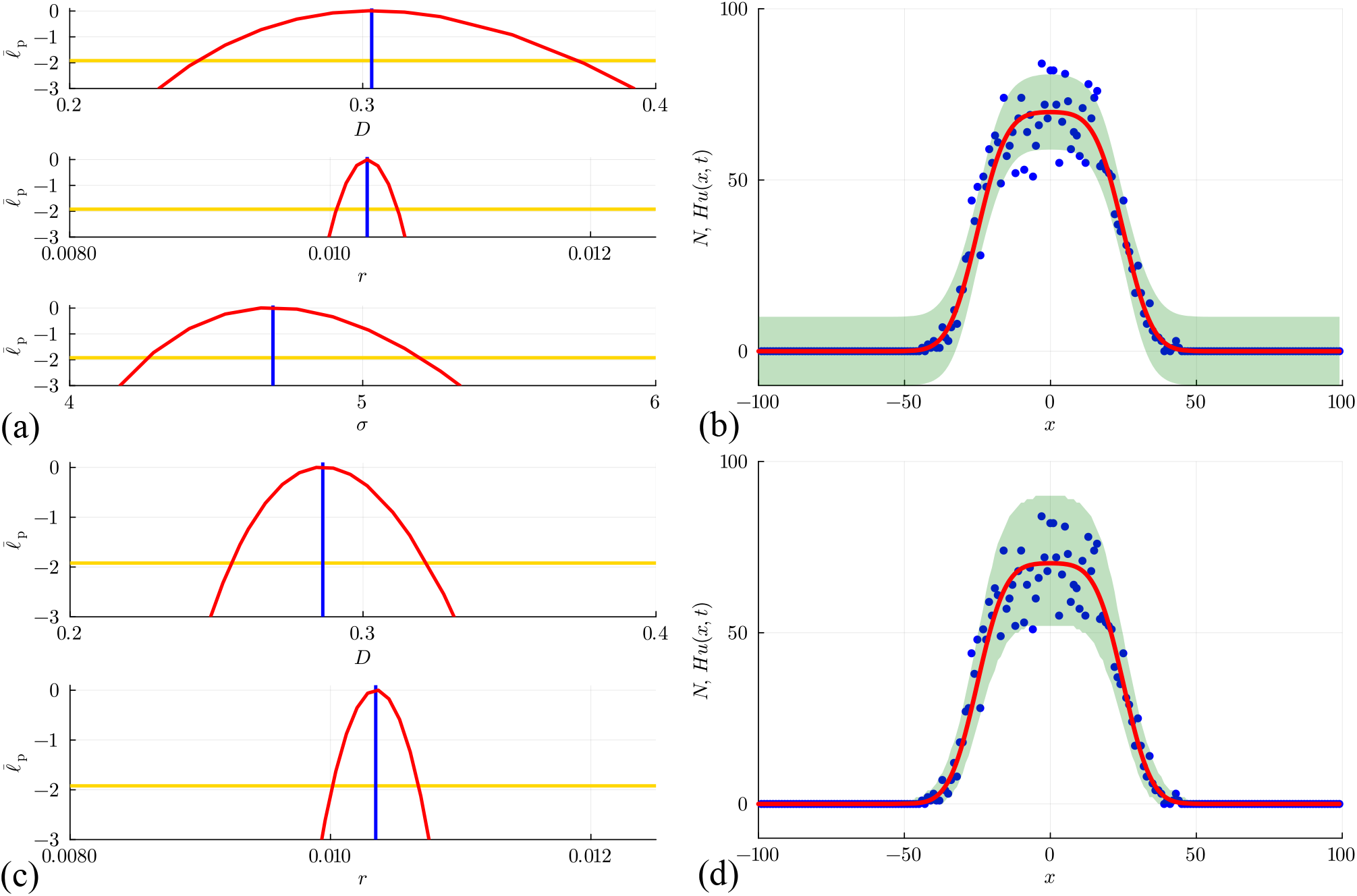
(a)–(b) Estimation, identifiability analysis and prediction under the additive Gaus _sian noise model. (a) Profiles for *D, r* and *σ* with 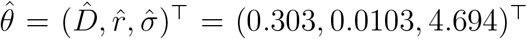_ superimposed (vertical blue lines). Each univariate profile gives an approximate 95% confi _dence interval: 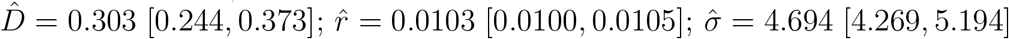_ generated by finding the values of the interest parameter where the profile intersects with the 95% threshold (yellow horizontal lines). (b) Prediction interval generated with the additive Gaussian noise model encloses 93.5% of the data. (c)–(d) Estimation, identifiability analysis and prediction under the Poisson noise model. (c) Profiles for *D* and *r* with 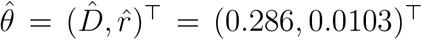. Each univariate profile gives an approximate 95% confi _dence interval: 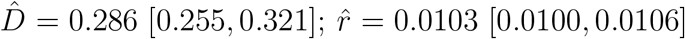_. (d) Prediction interval generated with the Poisson noise model encloses 99.0% of the data. Prediction intervals in (b) and (d) are generated using Laplace’s approximation.

Results in Figure 9(c)–(d) repeat the same estimation, identifiability and prediction steps under the Poisson noise model, leading to 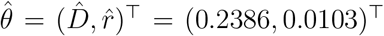, which is close to the expected result with each univariate profile likelihood function indicating that both parameters are practically identifiable. The prediction interval in Figure 9(d) reflects the positive count data more accurately than the additive Gaussian noise model approach, enclosing 99.0% of the data.

## 4 Interacting random walk models: Crowding effects

All results so far consider a classical non-interacting random walk model, where there is no limit to the number of agents that can occupy a lattice site. We now consider additional results for a very different model where individuals interact in a very simple way that can be taken as a simple model of crowding. These models are closely related to exclusion processes [33], where lattice site occupancy is limited to one agent per site. These models are often used in cell biology applications where cell-to-cell crowding effects are known to be very important [29, 36, 37, 82–89]. Similar interacting random walk models are also used in the context of ecological modelling applications [90–94]. The interacting random walk model is implemented on the same two-dimensional square lattice and we use the same random sequential update method [33]. During the updating algorithm, each selected agent is given the opportunity to move with probability *P* ∈ [0, 1], and the target site is chosen with directional bias so that a motile agent at site (*i, j*) will attempt to step to site (*i, j ±* 1) with probability 1*/*4, and to sites (*i±*1, *j*) with probability (1*±ρ*)*/*4. Any potential motility event that would place an agent on an occupied site is aborted. Once the *Q* potential motility events have been assessed, we then use the same random sequential update method to simulate agent proliferation. Each selected agent is given the opportunity to proliferate with probability *R* ∈ [0, 1]. A proliferative agent at site (*i, j*) will place a daughter agent at sites (*i ±* 1, *j*) and (*i, j ±* 1) with equal probability 1*/*4. Any attempted proliferation event that would place a daughter agent on an occupied site is aborted.

We now consider a suite of discrete simulations using this interacting random walk model on lattice with *H* = 50 and *W* = 200. As before, zero flux boundary conditions are applied along all boundaries, and simulations are initialised by randomly occupying each site within the region where |*x*| ≤ 25 with probability *U* = 0.5, leading to the distribution of agents shown in Figure 10(a). A single simulation is performed by allowing the system to evolve over 100 time steps with *P* = 1, *ρ* = *R* = 0 with the output of the simulation leading to a population that spreads symmetrically in the positive and negative *x*-directions, as shown in Figure 10(b). This initial and final distribution of agents is then summarised by plotting *N*_*i*_ at *t* = 0 and *t* = 100 in Figure 10(c)–(d), respectively. A key feature of these simulations is that *N*_*i*_ ≤ *H* owing to exclusion. A second simulation with *P* = 1, *ρ* = 0.5 and *R* = 0 is summarised in Figure 10(e)–(h) where we see the impact of the biased motion and exclusion. The spatial distribution of agents at *t* = 100 is asymmetric since motion in the positive *x*-direction is restricted by other agents in the population. The shape of the resulting distribution is fundamentally different to the biased simulation for the non-interacting random walk model shown previously in Figure 6. A third simulation with *P* = 1, *ρ* = 0 and *R* = 0.01 is summarised in Figure 10(i)–(l). This shows the impact of the growth and exclusion since the distribution of agents at *t* = 100 is restricted by exclusion and the total number of agents per column can be no greater than *H* = 50 in this case. The shape of the resulting distribution is fundamentally different to the non-interacting simulation with growth shown previously in Figure 8 where in the former case the number of agents per column is unlimited, whereas here exclusion enforces the condition 0 ≤ *N*_*i*_ ≤ *H*.

**Figure 10:**
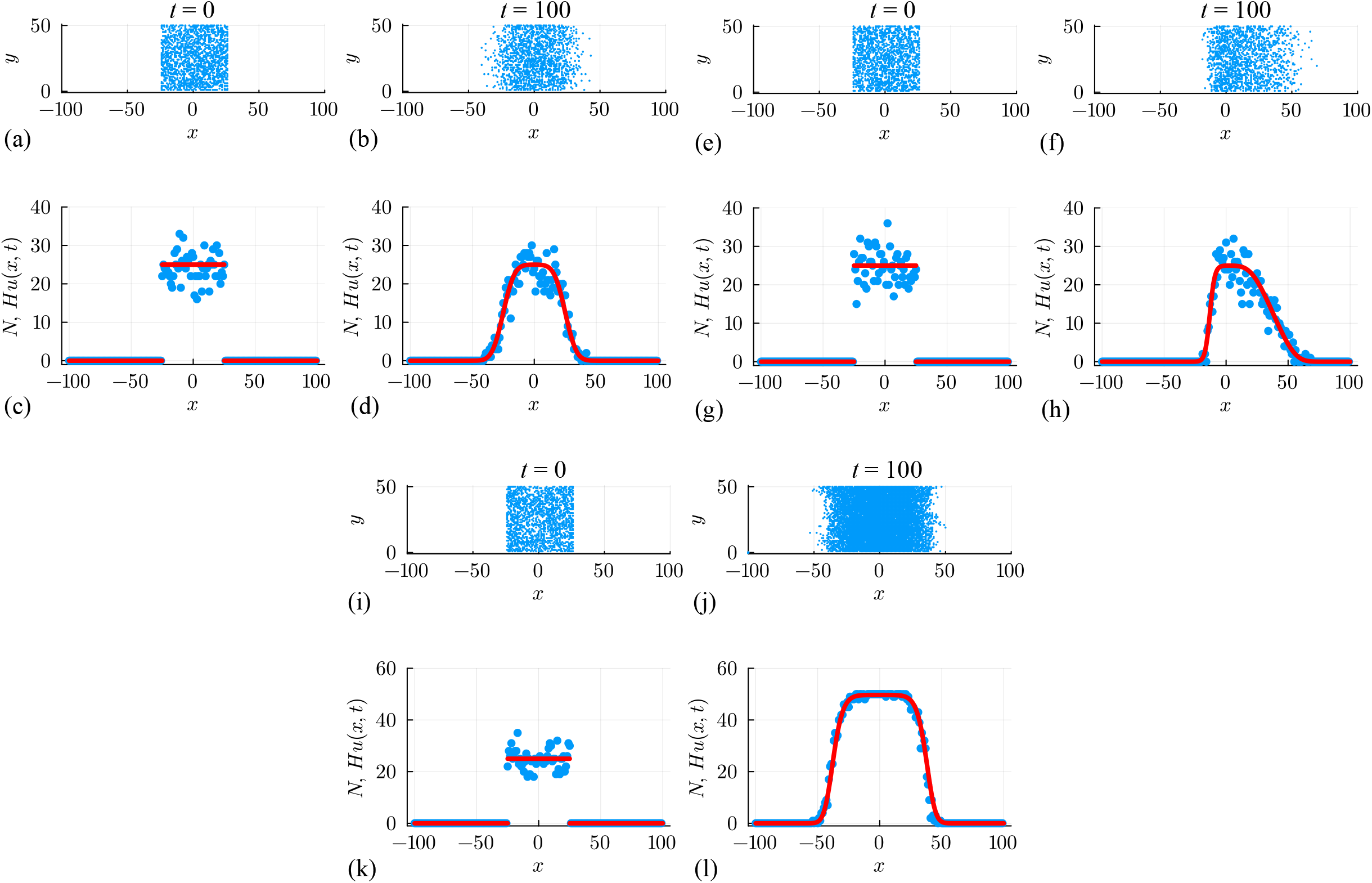
A suite of interacting random walk simulations on a square lattice with Δ = *τ* = 1, *H* = 50 and *L* = 100. Simulation results in (a)–(d), (e)–(h) and (i)–(l) correspond to: (i) an unbiased random walk (*P, ρ,R*)⊤ = (1, 0, 0)⊤, (ii) a biased random walk (*P, ρ,R*)⊤ = (1, 0.5, 0)⊤; and, (iii) an unbiased random walk with growth (*P, ρ,R*)⊤ = (1, 0, 0.01)⊤. In each case, the initial placement of agents in (a),(e),(i) involves all sites with |*x*| *≤* 25 being occupied by a single agent, at most, with probability *U* = 0.5, and the resulting arrangement of agents in (b),(f),(j) shows the resulting occupancy of sites at *t* = 100. Noisy count data (blue dots) in (c)–(d), (g)–(h), (k)–(l) is superimposed with *Hu*(*x, t*), where *u*(*x, t*) is the solution of Equation (20) with *D* = *P/*4, *v* = *Pρ/*2 and *r* = *R* (red curves).

We quantify the interacting random walk simulations in a similar way as before by computing *N*_*i*_, noting that now we have the restriction that 0 ≤ *N*_*i*_ ≤ *H*. The mean-field description can be obtained using the same conservation approach, giving [21]

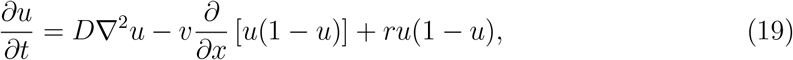

where 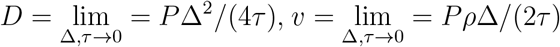 and 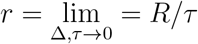, with *u*(*x, y, t*) ≥ 0 denoting the average density of agents at location (*x, y*) and time *t*. For simulations that do not involve macroscopic density gradients in the vertical direction the PDE simplifies to

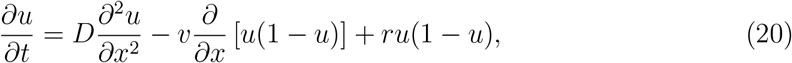

for the density *u*(*x, t*) ≥ 0. For simulations in Figure 10(a)–(d) with *ρ* = *R* = 0, this PDE relaxes to the linear diffusion equation and we can work with the same exact solution used previously, Equation (5). For simulations in Figure 10(e)–(l) with *ρ* ≠ 0 and/or *R* ≥ 0, we solve Equation (20) using a standard method-of-lines numerical approach, with details outlined in the Appendix. Numerical solutions of this nonlinear PDE model are superimposed on distributions of *N*_*i*_in Figure 10(g)–(h) and Figure 10(k)–(l), where we see that the solution of the nonlinear PDE model captures the mean trends in the simulation data quite well, but without capturing stochastic fluctuations.

Results in Figure 11(a)-(b) deal with the simulation data from Figure 10(a)-(d) for the interacting random walk model without bias or growth. Univariate profiles are superimposed _with 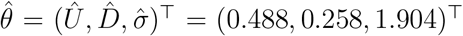_ are shown, together with the resulting prediction intervals under the standard assumption that *N*_*i*_| *θ* = 𝒩 (*Hu*(*x*_*i*_, *t*), *σ*^2^). In summary, we see that the MLE point estimate is close to the expected result, and the profiles indicate that the parameters are well identified by the data since we have well-defined, relatively narrow confidence intervals. As with previous implementations of the additive Gaussian noise model, the prediction interval encloses the stochastic data reasonably well, but leads to non-physical predictions of negative counts.

**Figure 11:**
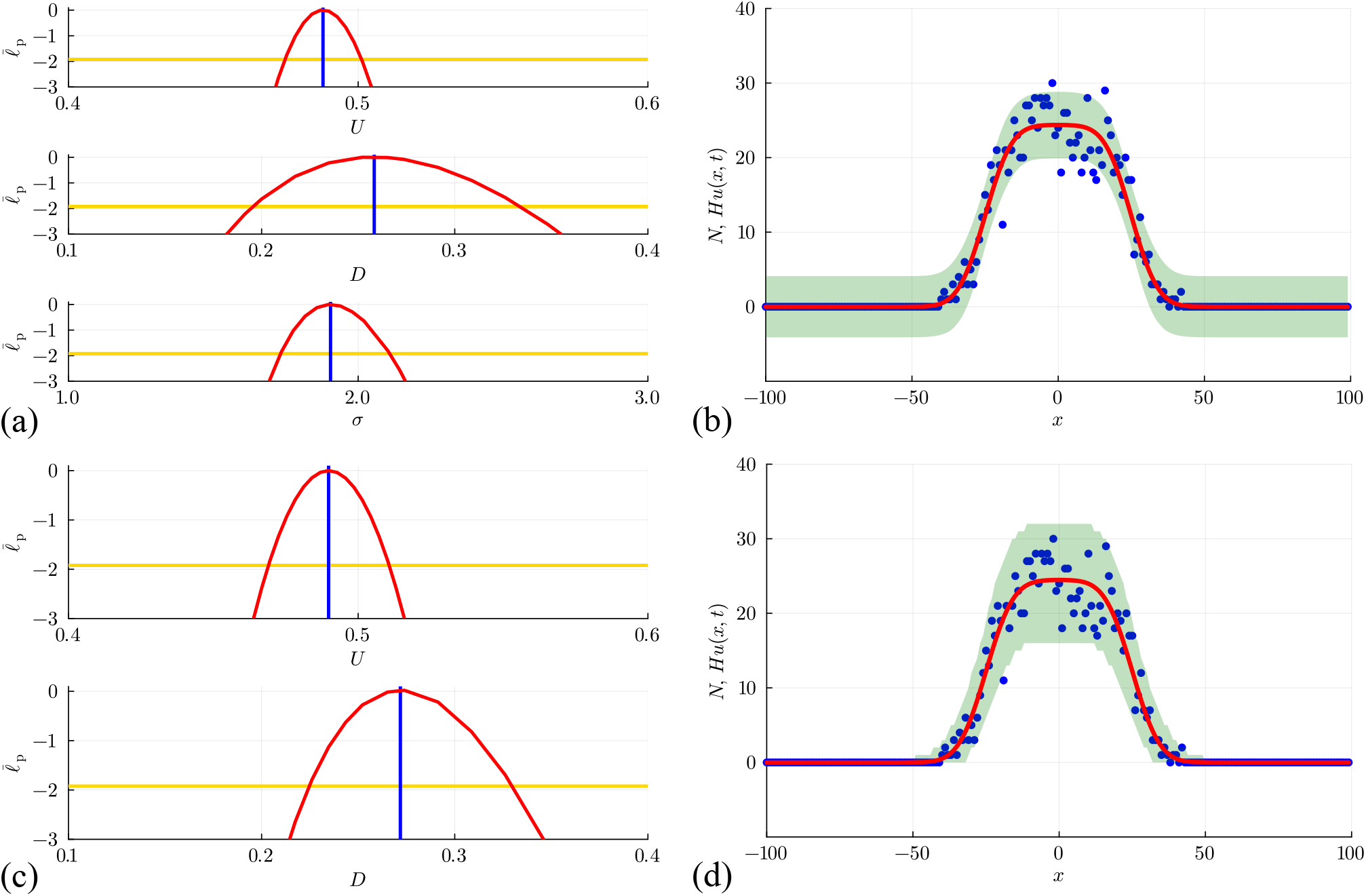
(a)–(b) Estimation, identifiability analysis and prediction under the ad _ditive Gaussian noise model. (a) Profiles for *U, D* and *σ* with 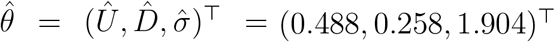_ superimposed (vertical blue lines). Each univariate profile gives an _approximate 95% confidence interval: *Û* = 0.488 [0.475, 0.501]; 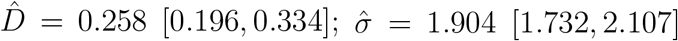_ generated by finding the values of the interest parameter where the profile intersects with the 95% threshold (yellow horizontal lines). (b) Prediction inter val generated with the additive Gaussian noise model encloses 95.5% of the data. (c)–(d) Estimation, identifiability analysis and prediction under the binomial noise model. (c) Pro _files for *U* and *D* with 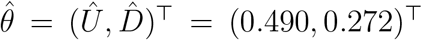_. Each univariate profile gives an _approximate 95% confidence interval: *Û* = 0.490 [0.469, 0.511];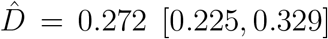_. (d) Prediction interval generated with the binomial noise model encloses 99.5% of the data. Prediction intervals in (b) and (d) are generated using Laplace’s approximation.

For the exclusion process model where each column has 0 ≤ *N*_*i*_ ≤ *H*, it is more natural to assume that the data *N*_*i*_ are drawn from a binomial distribution with *H* trials and success probability *u*(*x*_*i*_, *t*). For count data from the *i*th column we have

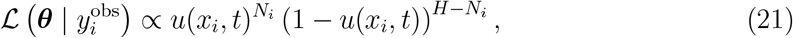

which, after invoking a standard independence assumption when considering a group of measurements at different location, *x*_*i*_ for *i* = 1, 2, 3, …, *I*, gives a loglikelihood function

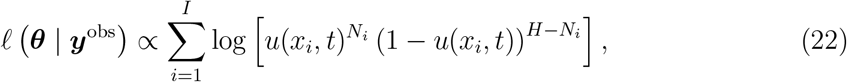

up to a constant. To evaluate this loglikelihood function we assume the proportionality constant is unity [39]. Working with this binomial loglikelihood function does not introduce any additional parameters. Results in Figure 11(c)-(d) show univariate profiles, superimposed with 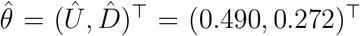, and the resulting prediction intervals. The MLE estimate is close to the expected result, and the profiles indicate that the parameters are well identified with well-defined, relatively narrow confidence intervals. The binomial loglikelihood function leads to a physically meaningful prediction interval that encloses 99.5% of the data, where the width of the prediction interval vanishes as *u*(*x, t*) → 0^+^ which precludes non-physical negative predictions.

To conclude we consider two final sets of results corresponding to discrete simulation data from the interacting random walk model with biased motion without growth in Figure 12, and for unbiased motion with growth in Figure 13. These results correspond to the simulation data associated with the snapshots in Figure 10(e)–(h), and Figure 10(i)–(l), respectively. In both sets of simulations, our count data is restricted to 0 ≤ *N*_*i*_ ≤ *H* for each *i* = 1, 2, 3, …, *I*, and the corresponding solution of Equation (20) must be obtained numerically. In both cases the main features of the noisy count data are captured quite well by the numerical solution of Equation (20). Unlike the non-interacting results with growth, shown in Figure 9 where *N*_*i*_is unbounded above, results in Figure 13 clearly show the impact of having one agent per site since by *t* = 100 the central columns near to *x* = 0 have *N*_*i*_ = *H* = 50 and the solution of the mean-field model appears to have approached a moving front. This is consistent with the observation that setting *v* = 0 in Equation (20) leads to the Fisher-Kolmogorov model which is well-known to have constant speed and constant shape travelling wave solutions [95].

**Figure 12:**
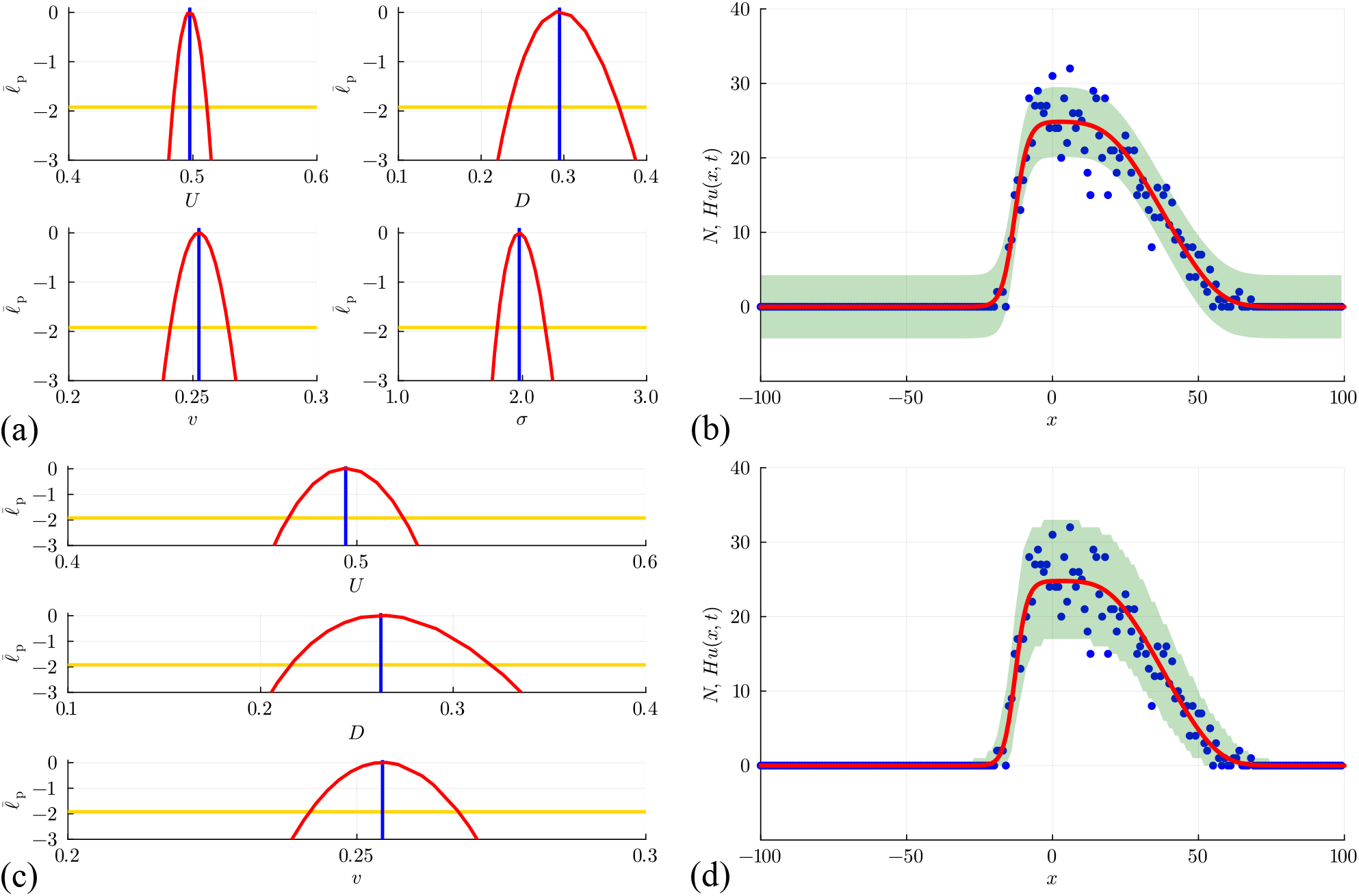
(a)–(b) Estimation, identifiability analysis and prediction under the addi _tive Gaussian noise model. (a) Profiles for *U, D, v* and *σ* with 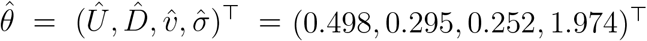_ superimposed (vertical blue lines). Each univariate profile gives _an approximate 95% confidence interval: *Û* = 0.498 [0.484, 0.511]; 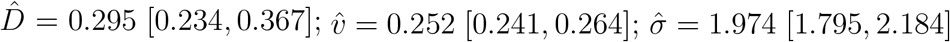_ generated by finding the values of the interest parameter where the profile intersects with the 95% threshold (yellow horizontal lines). (b) Prediction interval generated with the additive Gaussian noise model encloses 94.5% of the data. (c)–(d) Estimation, identifiability analysis and prediction under the binomial _noise model. (c) Profiles for *U, D* and *v* with 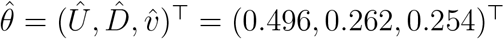_. Each _univariate profile gives an approximate 95% confidence interval: *Û*_ = 0.496 [0.476, 0.516]; 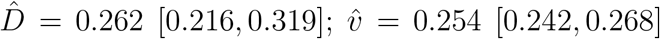. (d) Prediction interval generated with the binomial noise model encloses 98.0% of the data. Prediction intervals in (b) and (d) are generated using Laplace’s approximation.

**Figure 13:**
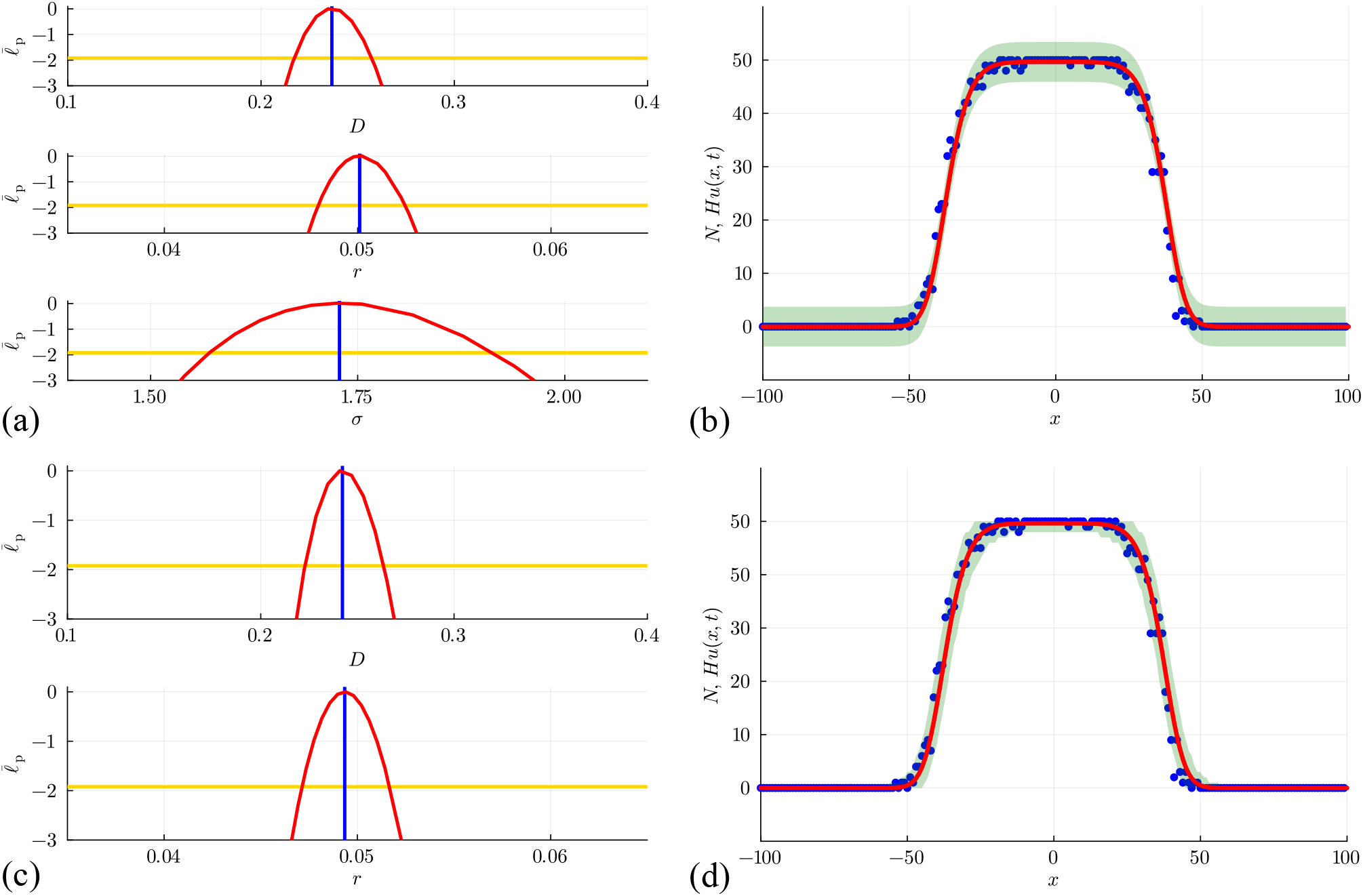
(a)–(b) Estimation, identifiability analysis and prediction under the additive _Gaussian noise model. (a) Profiles for *D, r* and *σ* with 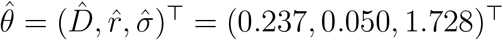_ superimposed (vertical blue lines). Each univariate profile gives an approximate 95% confi _dence interval: 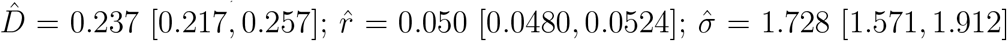_ generated by finding the values of the interest parameter where the profile intersects with the 95% threshold (yellow horizontal lines). (b) Prediction interval generated with the additive Gaussian noise model encloses 96.0% of the data. (c)–(d) Estimation, identifiability analysis and prediction under the binomial model. (c) Profiles for *D* and *r* with 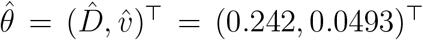. Each univariate profile gives an approximate 95% confi _dence interval: 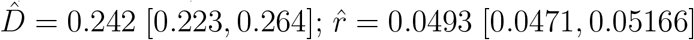_. (d) Prediction interval generated with the binomial noise model encloses 99.5% of the data. Prediction intervals in (b) and (d) are generated using Laplace’s approximation.

Figures 12(a)–(b) and 13(a)–(b) illustrate the outcome of likelihood-based estimation, identifiability, and prediction analysis under the standard assumption that count data are normally distributed about the solution of the mean-field model, *Hu*(*x, t*). While all parameters are identifiable, and prediction intervals enclose most of the stochastic data, both prediction intervals lead to non-physical outcomes. In particular, the prediction interval in Figure 12(b) indicates that negative counts are enclosed by the prediction interval, whereas the prediction interval in Figure 13(b) indicates that negative counts, and counts that exceed one agent per site, are enclosed by the prediction interval.

Figures 12(c)–(d) and 13(c)–(d) illustrate the outcome of likelihood-based estimation, identifiability and prediction analysis under the assumption that count data are binomially distributed with mean *Hu*(*x, t*). Under this noise model all parameters are identifiable, prediction intervals enclose most of the stochastic data, and both prediction intervals avoid non-physical outcomes.

## 5 Case study: Scratch assay

We conclude this review with a practical example. Before proceeding it is useful to note that all simulation and inference results so far are non-dimensional in the sense that we have set Δ = *τ* = 1 in all discrete simulations. This means that our estimates of *U, D, v*, and *r* are also non-dimensional. These non-dimensional simulations and parameter estimates can be re-dimensioned, as appropriate, to match a particular physical application by re-scaling Δ and *τ* with relevant length and time scales, respectively [34].

In this Section we will deal with a dimensional experimental data set, which means that we will work with dimensional quantities to match the appropriate time and length scales of a particular experiment, in this case a scratch assay [96–98] reported by Jin et al. [38]. Images of the experiment we consider are given in Figure 14(a)–(b) [38]. These images show the monolayer of cells just after a scratch has been made in Figure 14(a). A second image shows how the scratched region becomes recolonised by the impact of combined migration and proliferation in Figure 14(b) after *t* = 48 hours. These experiments use a human prostate cancer cell line, where individual cells have a diameter of approximately 20-25 *µ*m [38]. For the purposes of this Case Study we take the cell diameter to be a constant, Δ = 24 *µ*m. The images in Figure 14(a)–(b) are 1950 *µ*m wide and 1430 *µ*m high. Jin et al. [38] quantified the experiments by placing a series of uniformly-spaced vertical lines across each image and counting the number of cells in each column of width 50 *µ*m. To make this consistent with our simulation approach we make the further minimal assumption that cells are approximately uniformly distributed within each column, which allows us to report count data in Figure 14(c) that is equivalent to counting cells within columns of width 24 *µ*m so that the count data are in a similar format as the count data used in Sections 3.1–3.3. This involves rescaling the count data reported by Jin et al. [38] by a factor of 24*/*50, taking care to work with nearest whole numbers to preserve the fact we are working with non-negative integer count data. A simple square packing argument indicates that these columns can hold no more than 60 cells of diameter 24 *µ*m. Together, these minimal assumptions allow us to work with Equation (20) with *v* = 0 since there is no obvious asymmetry in the way that cells recolonise the scratched region of the experiments. This corresponds to working with the Fisher-Kolmogorov model [95, 99] as a surrogate process model, together with a binomial measurement error model to relate the noisy experimental count data to the solution of the surrogate model.

**Figure 14:**
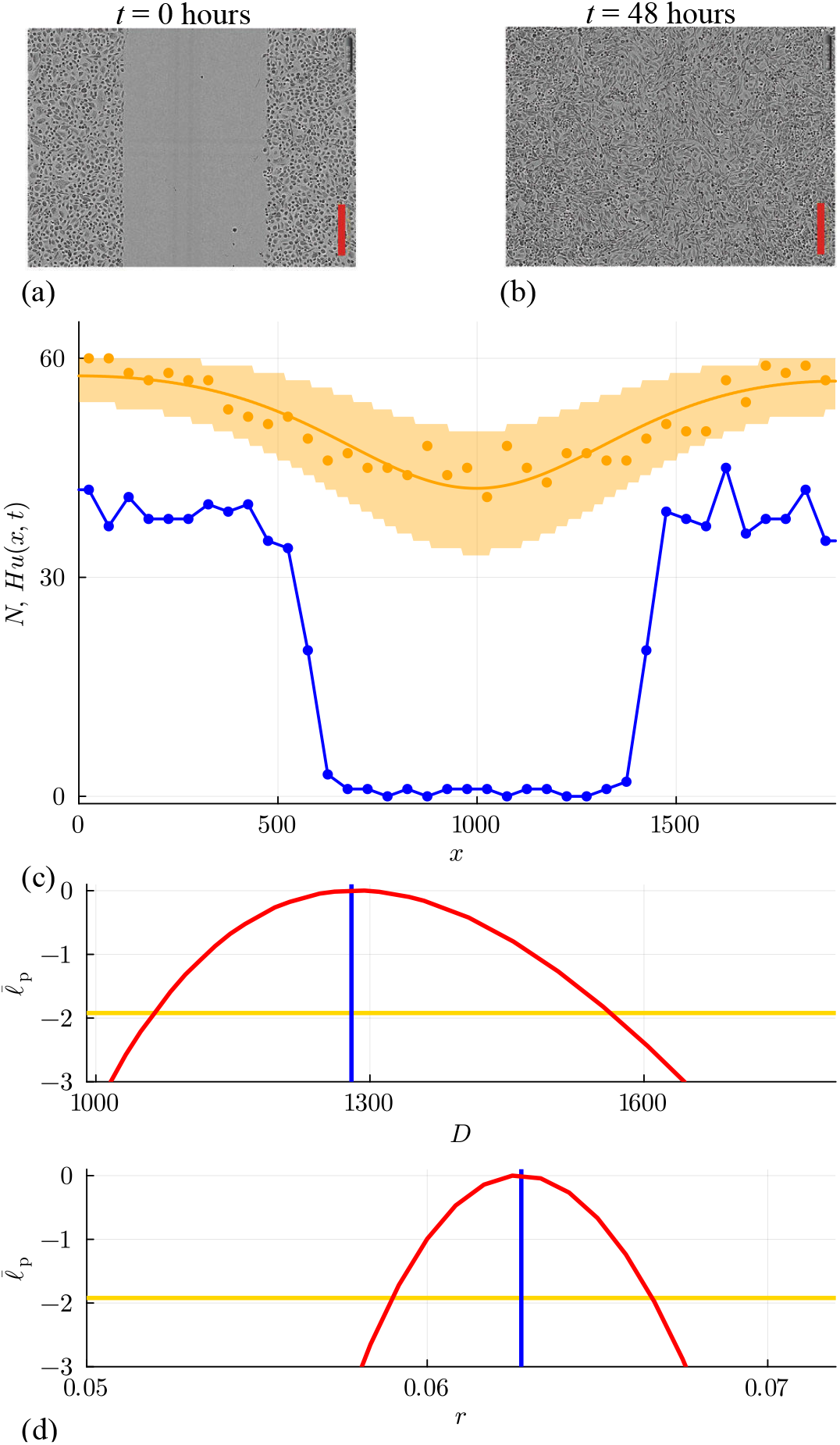
(a)–(b) Images of scratch assays reported by Jin et al. [38] at *t* = 0 and *t* = 48 hours, respectively. The red scale bar in each image corresponds to 300 *µ*m. (c) Count data at *t* = 0 (blue dots) and *t* = 48 hours (orange dots), together with the MLE solution of Equa _tion (20) with *v* = 0 and 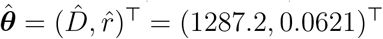_. The prediction interval in (c) for the data at *t* = 48 hours (shaded orange), generated using Laplace’s approximation, encloses 100% of the data. (d) Univariate profile likelihood functions for *D* (upper) and *r* (lower) superimposed with the MLE (blue vertical lines). Each univariate profile gives an approxi _mate 95% confidence interval: 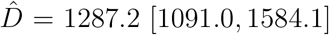_ and 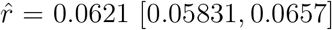 generated by finding the values of the interest parameter where the profile intersects with the 95% threshold (yellow horizontal lines).

Data in Figure 14(c) show the count data at *t* = 0 and *t* = 48 hours. Following Jin et al. [38] we linearly interpolate data at *t* = 0 to obtain the initial condition, which allows us to solve Equation (20) numerically to model the expected cell density profile at *t* = 48 hours. Using the binomial noise model, we obtain 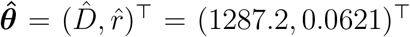, where the dimensional units of the cell diffusivity is *µ*m^2^/hour, and the dimensional units of the cell growth rate is /hour. These parameter values can be simulated using a discrete model with dimensional lattice size Δ = 24 *µ*m (representing one cell diameter). Recall that the macroscopic parameters *D* and *r* are related to the microscopic parameters via *D* = *P* Δ^2^*/*(4*τ*) and *r* = *R/τ*, where *P* and *R* are required to be in the interval [0, 1] as they represent probabilities. This imposes an upper limit on the choice of time step *τ* of 0.112 hours. For example, if we chose *τ* = 0.1 hours, this would lead to a movement probability per time step of *P* = 0.894 and proliferation probability per time step of *R* = 0.00621. These choices are not unique since we can choose a smaller value of the time step and simply re-scale our values of *P* and *R* to match our estimates of *D* and *r*.

Using automatic differentiation to calculate the Hessian matrix at the MLE, we invoke Laplace’s approximation to obtain the prediction interval in Figure 14(c) which encloses 100% of the data at *t* = 48 hours. As with the synthetic data examples, we see that the binomial noise model leads to bounded prediction intervals that faithfully captures the count data. Univariate profile likelihood functions in Figure 14(d) confirm that both parameters are identified by the data.

This case study provides a quantitative illustration of how we can interpret data from a cell biology experiment in a random walk framework using a surrogate PDE model. The original data presented by Jin et al. [38] involved reporting count data at *t* = 0, 12, 24, 36 and 48 hours. For the sake of simplicity, and to be consistent with the remainder of this review, here we work with data at the final time point only, *t* = 48 hours. It is straightforward, however, to incorporate the intermediate time points into our estimation and we will explain how to achieve this in Section 6.

We conclude the discussion of this Case Study by noting that interpreting this experimental data with a simple interacting random walk model and corresponding mean-field surrogate PDE description has limitations as there are many modelling assumptions that could be explored and relaxed. For the sake of clarity, here we interpret this experiment using data at a single time point which rules out the possibility of identifying time-dependent parameters. This is noteworthy because previous studies that examine this experimental data have used measurements at several time points and concluded that working with time-dependent parameters, *D*(*t*) and *r*(*t*), captures the experimental data across all time points more accurately than making the standard assumption that *D* and *r* are constants [39, 100]. In Section 6 we explain how to deal with data at multiple time points, however here we have not explored the possibility of time-dependent parameters, nor have we addressed more general questions about model selection. Nonetheless, despite working with a parsimonious model this case study illustrates how the ideas developed in Sections 3.1–3.4 can be applied to experimental data to give meaningful and interpretable mechanistic insights. More generally, when dealing with real biological data we are certain to face questions of model selection, and we recommend that parameter identifiability analysis ought to be combined with model selection analysis by prioritising identifiable models [101, 102].

## 6 Conclusions and future work

In this review, we have surveyed and implemented a range of options for parameter inference, parameter identifiability analysis, and model prediction for a suite of lattice-based random walk models that can be used to study collective motion in biological applications. After an initial exploration of a simple ABC rejection approach relying only on a high-fidelity but computationally expensive stochastic model, we primarily focused on using low-fidelity, computationally inexpensive surrogate PDE models of the form,

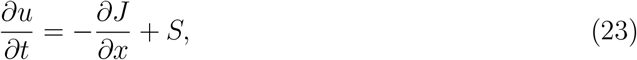

where *u*(*x, t*) ≥ 0 is the macroscopic mean agent density at position *x* and time *t, J* is the macroscopic agent flux, and *S* is the source term representing macroscopic agent proliferation [20, 21, 30–32].

We have illustrated how solutions of this surrogate PDE model, both exact and numerical, can be used for parameter estimation, identifiability analysis, and model prediction, within both likelihood-based sampling approaches and likelihood-based optimisation approaches. A particular focus of this review is to construct prediction intervals by combining various process models relating to details of the stochastic model, along with various noise models that relate the solution of the PDE to noisy observed data via a specified probability distribution [49]. A recurring result has been to show that the standard choice of working with an additive Gaussian noise model can lead to non-physical prediction intervals. In response, We illustrated how the use of other noise models (such as binomial and Poisson noise) more faithfully reflect the data-generation process, and can more accurately capture variability in the observed data.

In summary, our synthetic data analysis investigated 12 distinct random walk models, both non-interacting and interacting, whose key features are summarised in terms of the associated macroscopic PDE model in Table 1. While the surrogate PDE models in Table 1 describe problems posed in a one-dimensional Cartesian coordinate system, the general methods and workflow can also be applied when considering the same stochastic models and associated PDE descriptions in different coordinate systems. This includes radially symmetric problems and two- or three-dimensional Cartesian problems [103] without the vertical symmetry that we assumed in the synthetic simulation data and in the experimental Case Study presented in Section 5.

To maintain a straightforward style of exposition, we focus on cases where data are collected at a single time point. As mentioned in Section 5, while focusing on a single time point is relevant in many situations, it is also straightforward to extend our workflow to deal with data collected at multiple time points and spatial locations. For example, a modified log-likelihood function

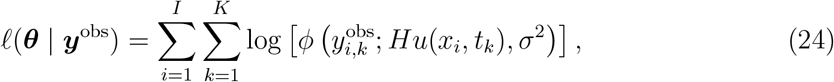

can be used to deal with data collected at *I* spatial locations, *x*_*i*_for *i* = 1, 2, 3, …, *I*, and at *K* temporal time points, *t*_*k*_ for *k* = 1, 2, 3, …, *K*, under a standard independence assumption. The key change here is that now the data vector *y*^obs^ is extended to include data collected at the *i*th spatial location and the *k*th time point, so that the length of this vector is *IK* and individual elements are denoted 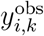. This extended form of the log-likelihood is written here for the standard case of working with an additive Gaussian noise model, and similar extensions apply for the other noise models considered in this review.

This review focuses on position-jump random walk models, where the continuum-limit surrogate model takes the form of a parabolic reaction-diffusion or reaction-advection-diffusion PDE [20, 21, 95]. We have not considered velocity-jump random walk models, where individual agents are associated with both a location and a velocity, and the continuum-limit surrogate model takes the form of a system of hyperbolic PDEs in a class known as transport equations [104]. Typically, non-interacting velocity-jump models lead to linear hyperbolic PDEs that are closely related to the telegraph equation [105–107], whereas interacting velocity jump models with crowding effects lead to nonlinear transport PDEs that resemble mathematical models of traffic flow [108]. The same framework and workflow we have outlined of collecting stochastic count data, representing the main trends in the data using a surrogate PDE model, and representing the stochastic data-generating process using an appropriate noise model can also be followed for velocity-jump models.

A key feature of the approaches for inference, identifiability analysis and model prediction in this review is to work with a computationally inexpensive surrogate model. While we have used standard mean-field approaches to arrive at surrogate PDE models [20, 21], there are alternative approaches for deriving appropriate surrogates. For example, methods of equation learning [100, 109–115] aim to *learn* the underlying structure of a differential equation model that gave rise to available observed data. These approaches have been used to generate surrogate ODE and PDE-based models and could be used instead of working with a mean-field PDE model. Other approaches include using a Gaussian process to construct a differential equation model from discrete data [116, 117] or using multidimensional linear regression as a computationally inexpensive option to represent any discrepancy between the solution of the mean-field PDE and averaged data from the discrete model [118]. It is also possible to make use of user-derived expertise to propose phenomenologically motivated differential equation models that can perform very well [119, 120].

Throughout this review, we have focused on identifiable problems where model parameters are reasonably well-identified by the data. A critical step in the workflow of parameter estimation, identifiability analysis, and model prediction is to quantify the extent to which the noisy data constrains our parameter estimates. Identifiability refers to whether a model’s parameters can be uniquely determined from observed data, and identifiability analysis involves evaluating how accurately and reliably these parameters can be estimated given the structure of the model and the data available [57]. Methods of identifiability analysis are typically classified in terms of whether they deal with *structural* or *practical* identifiability [25, 26, 50]. Structural identifiability analysis examines the question of whether different parameter values generate different probability distributions of the observable variables [58, 59, 121]. Structural identifiability is concerned with analysing the structure of the mathematical model, and is often assessed using software that use Lie derivatives to generate a system of input– output equations and examining its solvability properties [122–124]. Practical identifiability analysis concerns the extent to which parameter values can be confidently estimated, given noisy, incomplete data. This usually involves fitting a mathematical model to data, and then exploring the extent to which the quality of the model fit changes as the parameters are varied. Practical identifiability analysis can be performed either locally near a given point, such as near the parameter values that provide the best model fit, or globally over the extended parameter space. A common tool for assessing practical identifiability is the profile likelihood [23, 24, 60], which is the main focus on this work. Our choice to restrict our attention to reasonably well-identified problems explains why we chose to treat the initial density *U* as a known quantity when exploring random walk models with growth, *R >* 0. Intermediate to long-term data generated with random walk models with growth typically lead to identifiability challenges with the initial density *U*. This challenge was also circumvented in the case study where we treated the data at *t* = 0 as known and did not attempt to estimate the initial state of the system.

Since we focus on identifiable problems, there are many open questions about how to approach estimation and prediction for random walk models in the face of non-identifiability. Potential approaches involving reparameterisation to deal with structural non-identifiability [122– 125], and model simplification or improved experimental design and data collection protocols to deal with practical non-identifiability [70].

In this review we illustrate the use of Gaussian, Poisson, and binomial observation models. However, other distributions may be more appropriate in other applications, for example the negative binomial distribution for overdispersed count data, or the gamma or log-normal distribution for continuous, non-negative data. Switching to a different observation model is straightforward within the workflow we have presented as it simply involves substituting a different likelihood function into the inference method, whether that is profile likelihood or a Bayesian inference method. Throughout, we have assumed that noise in the observations at different locations is independent, but in the stochastic model (and in reality) fluctuations from the mean field at neighbouring locations will be correlated. Capturing non-independent noise is more challenging, and not covered by the approaches outlined here.

## Data Accessibility

Julia implementations within Jupyter notebooks for all computations are available on GitHub https://github.com/ProfMJSimpson/RandomWalkInference.

## Funding

This work is partly supported by the Australian Research Council (DP230100025, CE230100001) and the Marsden Fund (24-UOC-020).

## Acknowledgements

We thank the MATRIX mathematical research institute for hosting a one-week residential workshop entitled “Parameter identifiability in mathematical biology” (September 2024) where initial work on this project took place. We thank two referees for their supportive comments.

## Appendix

We obtain numerical solutions of Equation (20) by discretising the domain −*L* ≤ *x* ≤ *L* using a uniform mesh, with mesh spacing *δ*. Each mesh point has position *x*_*i*_for *i* = 1, 2, 3, …, *I* and we write the solution *u*(*x*_*i*_, *t*) as *u*_*i*_. Discretising spatial derivatives using standard central difference approximations leads to

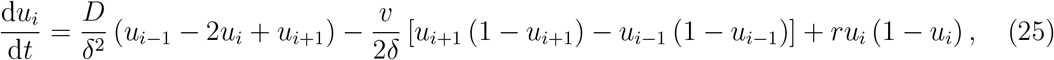

for *i* = 2, 3, 4, …, *I* − 1, together with boundary conditions *u*_1_ = *u*_*I*_ = 0. We integrated this system of ODEs through time using Julia’s DifferentialEquations.jl [126] using standard tolerance choices and automatic time stepping with Heun’s method. All numerical results presented in this review involve *δ* = 0.5, and additional results (not shown) indicate that this choice leads to grid-independent results for the problems that we consider.

## Notes

https://github.com/ProfMJSimpson/RandomWalkInference

### Competing Interest Statement

The authors have declared no competing interest.

### Summary of Updates

We have made a number of small changes, incluing adding some text to give a self-contained introduction to the concept of parameter identifiability, and to give a more detailed explanation about the use of Wilks' theorem to establish confidence sets and confidence regions.

